# On effect of chloroform on electrical activity of proteinoids

**DOI:** 10.1101/2023.12.29.573657

**Authors:** Panagiotis Mougkogiannis, Andrew Adamatzky

## Abstract

Proteinoids, or thermal proteins, produce hollow microspheres in aqueous solution. Ensembles of the microspheres produce endogenous spikes of electrical activity, similar to that of neurons. To make a first step towards evaluation of the mechanisms of such electrical behaviour we decided to expose proteinoids to chloroform. We found that while chloroform does not inhibit the electrical oscillations of proteinoids it causes substantial changes in the patterns of electrical activity. Namely, incremental chloroform exposure strongly affect proteinoid microsphere electrical activity across multiple metrics. As chloroform levels rise, the spike potential drops from 0.9 mV under control conditions to 0.1 mV at 25 mg/mL. This progressive spike potential decrease suggests chloroform suppresses proteinoid electrical activity. The time between spikes, the interspike period, follows a similar pattern. Minimal chloroform exposure does not change the average inter-spike period, while higher exposures do. It drops from 23.2 min under control experiments to 3.8 min at 25 mg/mL chloroform, indicating increased frequency of the electrical activity. These findings might leads to deeper understanding of the electrical activity of proteinoids and their potential application in the domain of bioelectronics.

## 1. Introduction

Thermal polymerisation of mixtures of amino acids results in the formation of dynamic oligopeptide structures — proteinoids — that tend to grow continuously [1, 2, 3]. These structures also assemble into larger aggregates at both microscopic and macroscopic scales [4, 5, 6, 7, 8]. Proteinoids, found within these structures, display intricate emergent chemistry such as imitating ion channels, regulating voltage, and responding to stimuli in a manner similar to living cells [9, 10, 11, 12]. Proteinoids are bio–inspired materials that can mimic the biophysical and self–organisational characteristics observed in living organisms [13, 14, 15, 16, 17, 1].

While previous research focused on morphological characteristics [2, 18, 19, 20], recent advances in electrochemical analysis provide critical insights into bio–electric activity [21]. Evidence suggests that proteinoid aggregates have significant self–propagating electrical currents with dynamic oscillations resembling neuronal potentials [22, 23, 24, 25].

Proteinoids exhibit a specific susceptibility to chemical stimuli that cause alterations in their morphology [26, 27, 28]. Changes in both the structural and electrostatic binding affect the tertiary conformations and likely the properties of ion permeability [29, 30].

Proteinoid microsphere produce spikes remarkably similar to the action potential [23, 24] (Fig. 1). The precise mechanisms underlying proteinoid excitability are not universally agreed upon. The physical conditions governing excitability are delineated in [31]. One theoretical model posits that excitability is contingent on the strength of coupling between basic and acidic proteinoids, which undergo abrupt changes over time due to material flow [8]. Additional models worth considering include the classical model involving ionic gradients established through channels [32] and the generation of membrane potential through the immobilisation of mobile ions. In this model, ions adhere to the surface of proteinoid microspheres, thereby generating a nonzero membrane potential [33]. To evaluate if excitability of the proteinoids relates to excitability of living substrates, we decided to conduct experiments with chloroform. Another rational for conductive experiments discussed presently, is to evaluate how close are proteinoid microspheres to living forms. Here we follow Claud Bernard’s hypothesis that sensitivity of anaesthetics differentiate living substrates from inanimate chemical forms [34].

**Figure 1:**
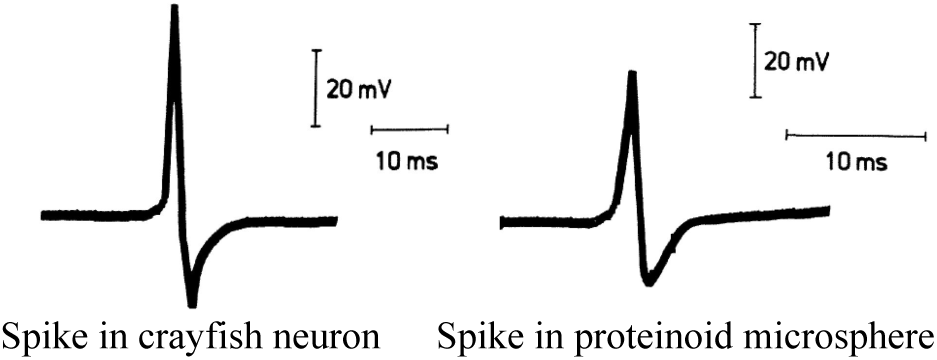
Action potentials from crayfish stretch receptor neuron and from proteinoid microsphere. Modified from [22].

Chloroform is a chemical compound employed as an anaesthetic, which effectively inhibits nerve signals and induces paralysis in certain parts of the mammals’ body [35, 36]. Chloroform acts as anaesthetic not only on mammals and their single cells [37] but also plants [38, 39, 40, 41] fungi [42], slime moulds [43] and bacteria [44].

Chloroform is believed to exert its effects by permeating cell membranes, according to researchers [45, 46]. These membranes possess channels that facilitate the passage of ions and messages. When chloroform becomes trapped in the membranes, it obstructs the channels, impeding the normal transmission of signals [47, 48].

Through the examination of chloroform’s impact on synthetic cell membranes, we can enhance our understanding of these phenomena [49, 50]. Membranes were synthesised utilising “proteinoids”, which are polypeptides composed of amino acids. The proteinoids have characteristics similar to cell membranes, forming small vesicles [51].

Efficient signal transmission is vital for the optimal functioning of nerve cells. Therefore, it is crucial to understand techniques for restoring conduction in order to maintain optimal health. In addition to its medical applications, studying membrane signal plasticity and adaptive mechanisms can also contribute to unconventional computing [52, 53]. Exploring proteinoid excitability phenomena [54, 23] could unlock exciting possibilities for innovative data processing methods that utilise the intricate ionic processes [55, 56] found in living organisms. More precisely, manipulating the restore/collapse spiking in tailored bio–polymer systems could offer a pathway for conducting multi–state logic operations. Phase transitions that are influenced by the environment can potentially be used to convert chemical signals into digital data, affecting the rhythm and appearance of the signals. Designing robust and adaptive bio–electronic systems involves cataloguing emergent identity resiliency factors. These systems are capable of self–recovering vital communication pathways, even in the face of disruption threats. Efforts to rescue delicate biological conduction phenomena have led to foundational understandings that ignite bio–inspired innovation in sensing, computing, and tissue engineering [57, 58]. By looking at signal stability from a different perspective, this research gains even more promise and urgency.

In this study, we explore the effects of varying chloroform concentrations on the dynamic electrochemical indicators in thermal proteinoid polymers. We measure shifts in oscillatory rhythms, periodicity, spike intensity, and duration across a range of exposures, from trace amounts to anaesthetic concentrations. The insights gained provide clarity on tuning ranges for sustained, regulated bio–electronic phenotypes in the presence of solvent disruption, thereby guiding the development of tunable signal–processing materials [59]. Proteinoids, comprised of thermal amino acid polymers, have been shown to self–assemble into cell– like microspheres able to exhibit lifelike phenomena. Yet, an enhanced understanding is still needed of the underlying structural dynamics that enable proteinoids’ biomimetic activities. Advanced computational modeling as shown for a small L–glutamic acid:L–arginine system (Fig. 2) can help reveal mechanisms related to spontaneous peptide aggregation, organisation, and conformational response in proteinoids by simulating their molecular behaviour.

**Figure 2:**
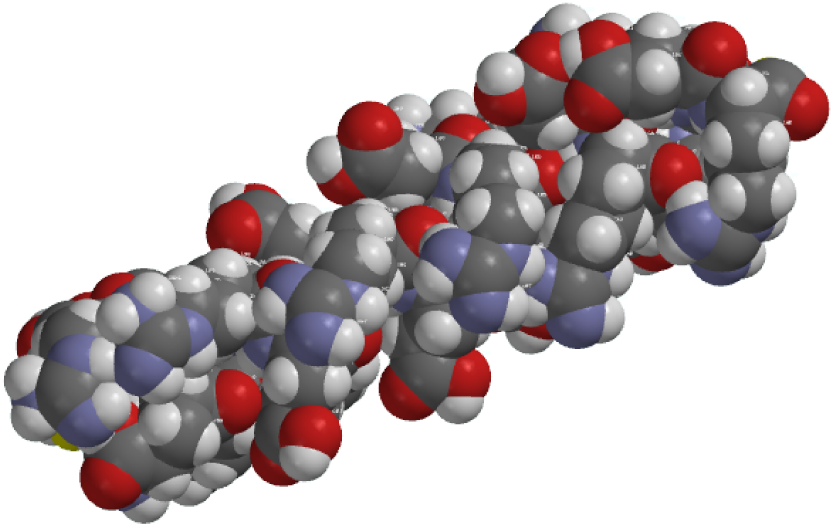
Predicted via molecular dynamics simulations, a space–filling model of a L-glutamic acid:L-arginine proteinoid ensemble consisting of eleven L–Glu:L–Arg peptide sequences. Blue denotes nitrogen, red represents oxygen, and grey represents carbon in the atomic colour scheme. Under the simulated environmental conditions, the total system minimum energy was −7995.08 kJ/mol. The random polymerization of glutamic acid and arginine during the thermal proteinoid synthesis imparts the self–assembled microsphere with a uniform distribution of constituent amino acids. The process by which oxygen moieties aggregate towards the outer surface is due to hydrophilic interactions with the aqueous solution phase. The AVOGADRO programme was utilised to conduct molecular dynamics simulations [60].

## 2. Methods and Materials

The amino acids L-Aspartic acid, L-Glutamic acid were procured from Sigma Aldrich, ensuring a purity level exceeding 98%. Upon procurement, the thermal polycondensation technique put forth by Mougkogiannis et al. [4] was implemented for the synthesis of proteinoids. This procedure comprised heating equimolar mixtures of the amino acids at 180°C, throughout a nitrogen atmosphere. The heating process was carried out for 30 minutes with consistent stirring. Post-heating, the proteinoids were subjected to lyophilization and stored at ambient temperature. For the experiments undertaken, chloroform was procured from Sigma Aldrich. It is listed under the CAS Number 67–66–3, possessing a molecular weight of 119.3 gr/mol. An FEI Quanta 650 microscope was used to capture the morphological attributes of proteinoids via Scanning Electron Microscopy (SEM). Notably, the gold sputter–coating procedure that is commonly employed in SEM imaging to enhance conductivity was skipped, underscoring the inherent conductive properties of proteinoids even under high–vacuum conditions Characteristics of I–V were documented with the employment of a Keithley 2450 sourcemeter. Platinum–iridium coated stainless–steel sub–dermal needle electrodes, procured from Spes Medica S.r.l., were implanted into the proteinoids allowing a nearly 10mm distance between each electrode pair. Moreover, the electrodes’ electrical activities were chronicled accurately by the use of a high–resolution data logger strapped with a 24–bit analog–to–digital converter – ADC–24 from Pico Technology, UK. The chronoamperometry measurements integral to our study were carried out using an Anapot EIS potentiostat.

The glass electrochemical cell used was a standard 5 mL vial (Fig. 3a). Platinum-iridium electrodes (Fig. 3b) served as the working and counter electrodes for applied voltage and current measurements. A Picolog ADC–24 (Fig. 3c) recorded voltage responses via the immersed electrodes. Custom software (Fig. 3e) controlled the Picolog unit and logged current data on a connected computer (Fig. 3d).

**Figure 3:**
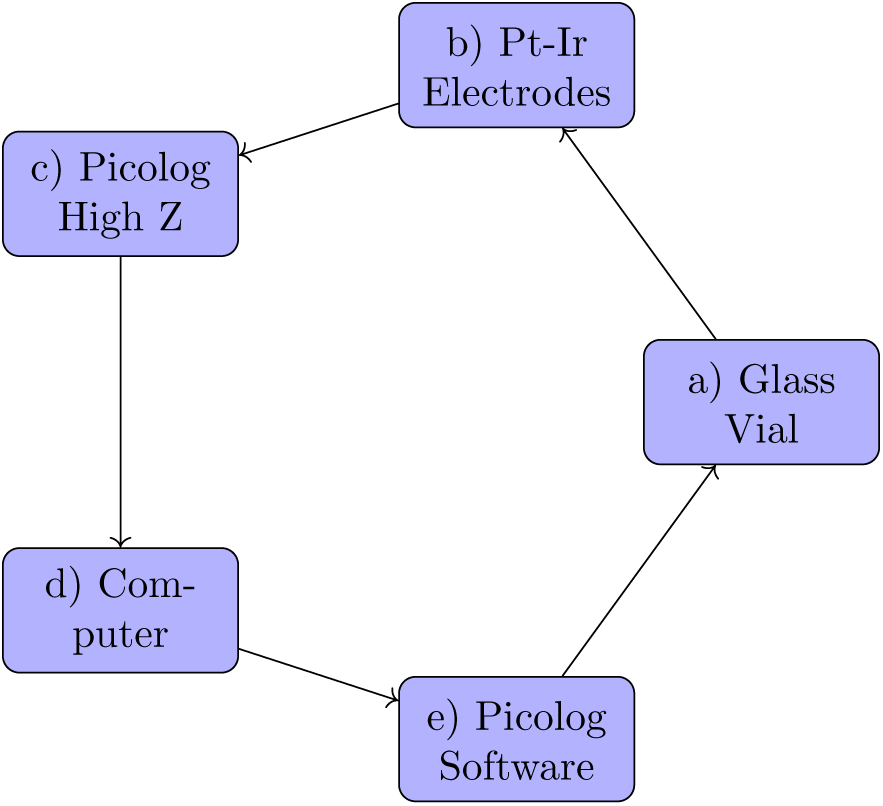
Key experimental equipment for voltage characterisation of proteinoids dynamics.

## 3. Results

Polymerised microspheres were subjected to step–wise chloroform exposure regimes and the resulting bio–electric responses were recorded to assess the impact of chloroform on proteinoid activity dynamics. The intrinsic spiking potentials of control proteinoid assemblies reflected baseline excitation characteristics. The addition of chloroform–soaked filter paper gradually increased the vapour phase concentrations in the container. To generate an escalating exposure series, 0.5×0.5 cm, 1×1 cm, and 3 ×3 cm filter sheets were saturated with chloroform and suspended over proteinoid solutions. Finally, direct chloroform addition into proteinoid samples at 25 mg/L investigated responsiveness to solvated phases.

This range encompasses anaesthetic concentrations for aquatic organisms, allowing examination of neurological excitation effects. Being denser than water, the chloroform likely partitions to lower layers, thereby inducing gradient diffusion. The systematic escalation in chloroform levels allows quantification of concentration–dependent modulatory impacts on proteinoid interspike intervals and spike morphologies. Contrasting control dynamics with responses across exposure regimes provides insights into intrinsic and externally–triggered influences on emergent bioelectric behaviours in this thermal protein polymer system.

Scanning electron microscopy elucidated both the baseline ultrastructure (and impact of solvent exposure on proteinoid morphologies, as exemplified in Fig. 4a. However, when exposed to chloroform, the broken microspheres developed bursting membranes and hollow interiors (Fig. 4b). This microscopy evidence supports the hypothesised swelling and permeation effects.

**Figure 4:**
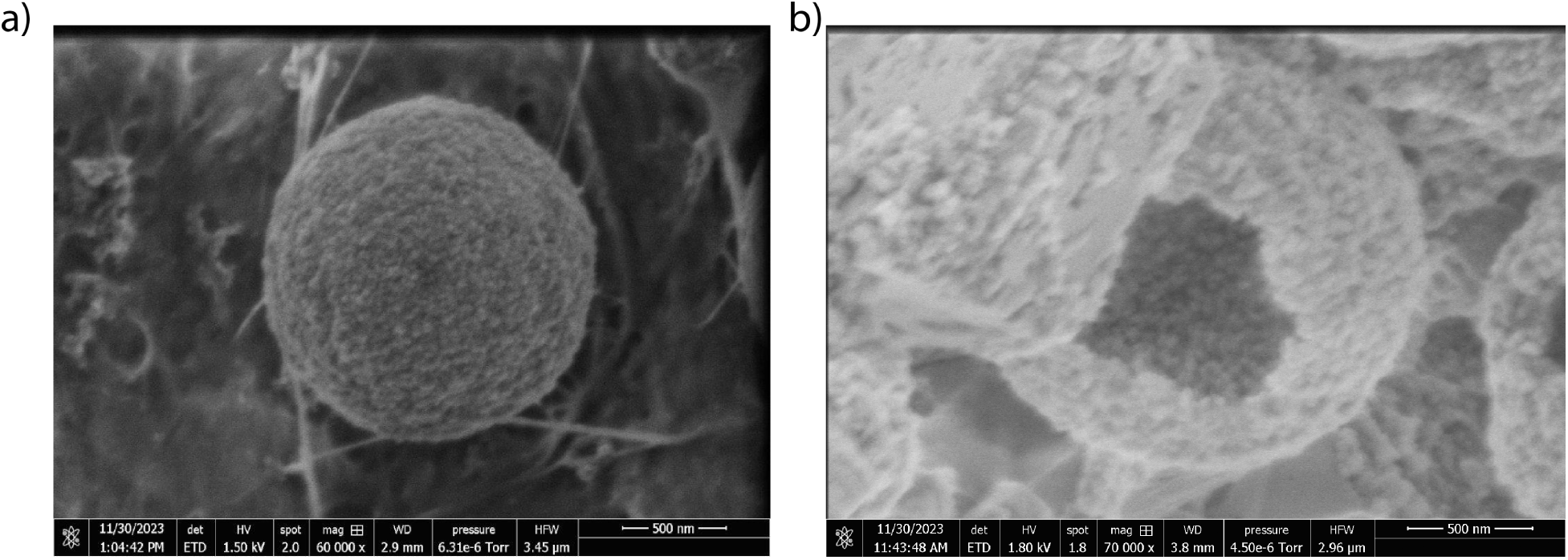
Scanning electron micrographs of L–glutamic acid:L–arginine (L-Glu:L-Arg) proteinoids microspheres (a) before and (b) after chloroform exposure. (a) Shows a representative intact single proteinoid particle with rough surface morphology and solid interior (scale bar 500 nm, 1.50 kV beam, x60,000 magnification). (b) Displays a hollow chloroform–treated microsphere with fractures enabling view of internal structure consisting of smaller nanosphere subunits (scale bar 500 nm, 1.80 kV beam, x70,000 magnification). Solvent ingress likely strains and ruptures the proteinoid membrane boundary, revealing the presence of embedded particulate layers inside. This supports a layered particle assembly formation mechanism, which gets disrupted upon solvent penetration.

### 3.1. Baseline Electrical Dynamics

According to spike analysis, the oscillations recurred with an average periodicity of 1392.86 s (Fig. 18 and Fig. 5) with amplitudes ranging from 0.36 mV to 1.91 mV (*µ*= 0.89 mV; *σ*=0.42 mV) Fig. 24.

**Figure 5:**
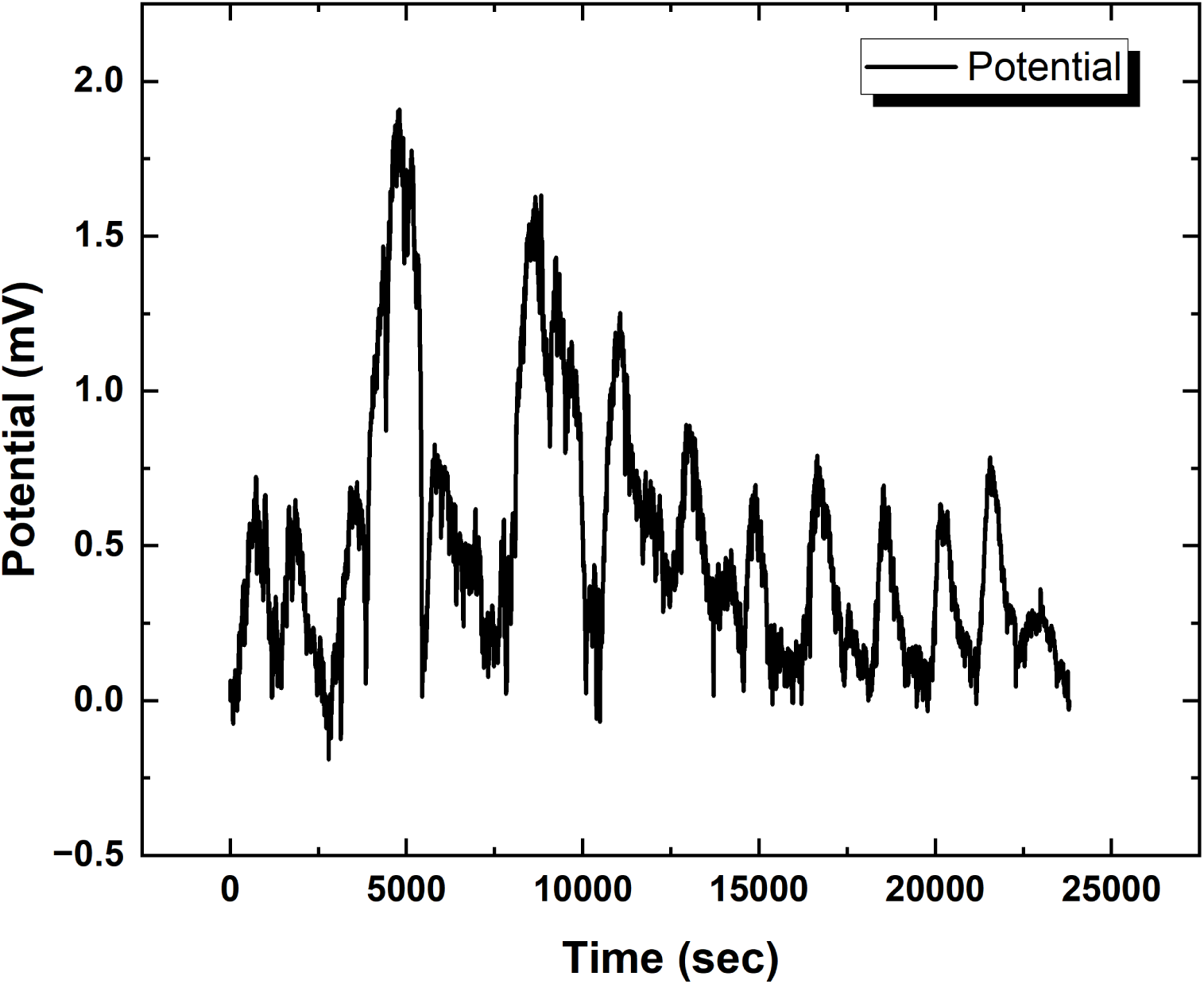
Proteinoid bioelectric activity after removing the baseline. The graph displays the temporal pattern of proteinoid spiking activity after applying a B–spline fit and removing the baseline signal. Through the extraction of slower field potential alterations, one can uncover and examine the dynamics of spike pattern. Spikes observed at around 5000 seconds and 7000 seconds have much higher densities in comparison to the intervals in between and subsequent to those timeframes. Modifications in spike characteristics such as intensity, breadth, or clustering due to external stimuli can provide insight into the adjustment of emergent excitation properties in these artificial polymers. This serves as the control sample showing baseline L–Glu:L–Arg proteinoid excitation dynamics without the effects of chloroform for comparison.

The amplitude intensity distribution was positively skewed (skewness = 1.37) and leptokurtic (kurtosis = 3.91), indicating a lengthy right tail of high amplitude spikes that deviated from the mean. This shows that the dynamics of proteinoid excitation entail occasional significant bursts mixed with more constant suppressed oscillations.

Control proteinoid electrical activity exhibits complexity and stochasticity, as seen in the signal shape variations in Fig. 18 across time and across repeated measurements. Profiling spike shape attributes further could provide insight into how microscale structural assembly events appear in the reported macroscale potential patterns.

Under control conditions in the absence of chloroform, the spiking activity of L–Glu:L–Arg proteinoid samples varied significantly (Fig. 19). Interspike intervals measured varied from 595.24 to 1952.38 s, with a mean of 1392.86*±*106.46 s (Fig. 19A,B). The distribution has a negative–skew Gaussian–like shape (skewness = −0.36) and a low kurtosis (1.95). The heterogeneous rhythm suggests that proteinoid oscillations develop from complex dynamics even when no external perturbations are present. Changes in these distributions in response to alterations such as the addition of chloroform may reveal environmental influences of proteinoid bioelectric behaviour. Identifying characteristics that modulate rhythmic spiking could provide insight into how to programme and improve interface with these synthetic mimics of excitable membranes.

### 3.2. Vapour Phase Chloroform Exposures

#### 3.2.1. Modulation of Proteinoid Spiking Upon Exposure to 0.5 cm × 0.5 cm Chloroform–Embedded Filters

Our analysis of Figs. 6 and 20 reveals intriguing findings regarding the response of proteinoids to the incorporation of chloroform in the vapour phase. We observe a reduction in spike amplitudes and a shift in inter–pulse intervals, indicating that the protein system regulates internal excitation dynamics when faced with external perturbation.

**Figure 6:**
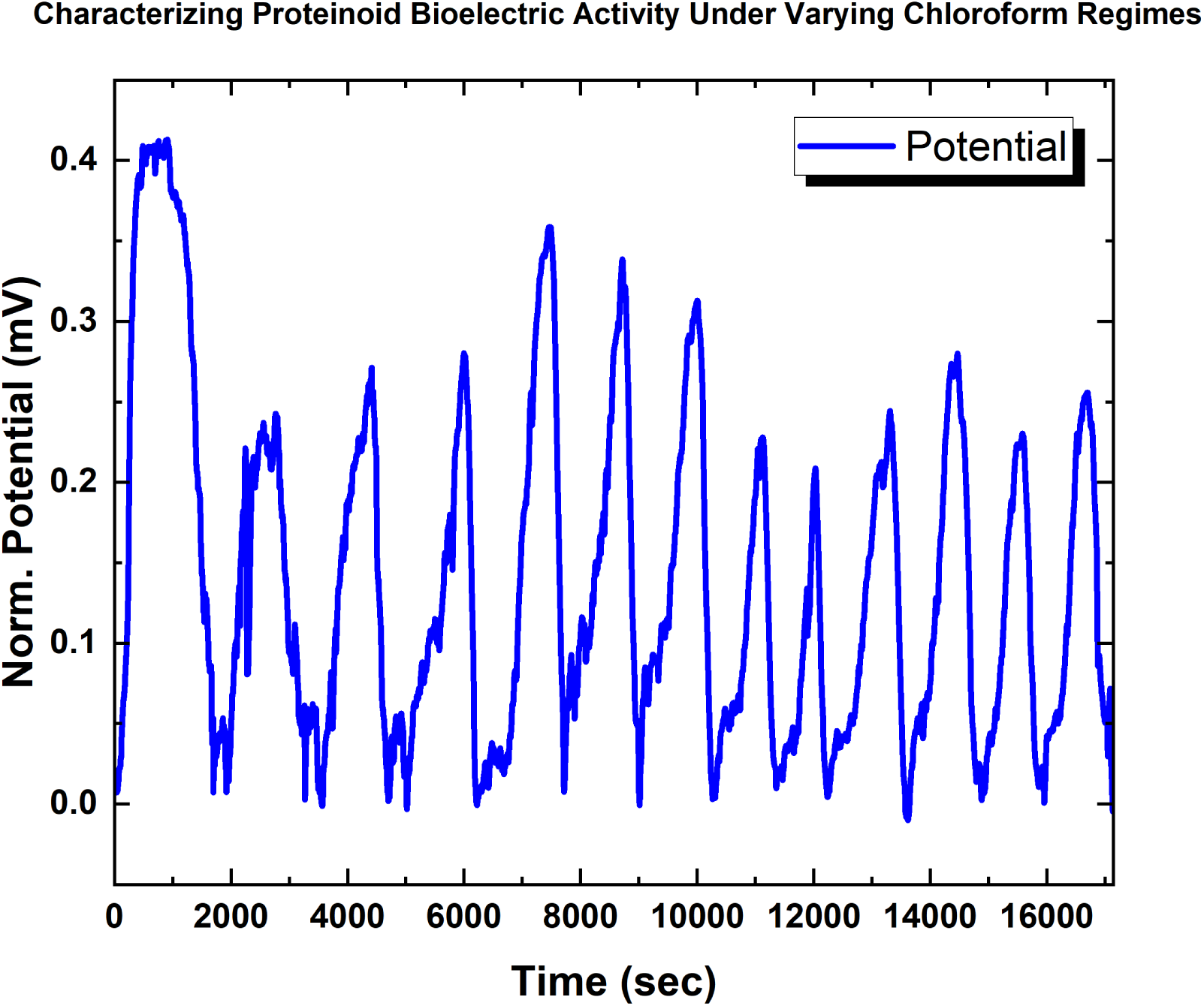
Proteinoid spiking amplitude dynamics with 0.5*×*0.5 cm^2^ chloroform filter paper exposure. When compared to greater baseline levels without chloroform, the timecourse indicates amplitude fluctuation over time (mean = 0.28 mV; max = 0.41 mV, min = 0.21 mV, *σ* = 0.06 mV). The distributions retain positive skew (0.87) but have lower kurtosis (2.83) compared to the control. Under limited chloroform, attenuation and confined amplitude ranges suggest dose–dependent regulation of excitation dynamics.

Notably, the distribution of amplitudes shows a decrease in variability, while maintaining a positive skewness. This suggests that the conductivity deviations are limited, yet the overall activity remains outward. Additionally, we observe shortened periods and accelerated rhythms, indicating the generation of actionable potentials to counter the disruptive effects of diffusive chemicals. In essence, we find that the proteinoid architectures exhibits self–stabilisation reflexes when confronted with low–level environmental interruptions.

The baseline mean spike potential of control proteinoid samples (Fig. 25) was 0.89*±*0.42 mV. The addition of a 0.5×0.5 cm^2^ chloroform source caused observable alterations in spiking activity. When exposed to filter– sourced vapours, the average spike potential was 0.281*±*0.016 mV. This is a 68% reduction in spike intensity when compared to the control spike intensity without chloroform. The significant potential attenuation supports feasible chloroform diffusion gradients reaching proximal proteinoid microspheres and incorporating into assemblies.

The continuous pattern of spiking activity, on the other hand, indicates preserved bioelectric functioning. Further incremental scaling of vapour phase chloroform via enlarged filter sheets or new sources may intensify adaptive dynamics dose–dependently. Comparing morphological and conductivity readouts across various exposure scopes may help to identify mechanisms that allow resonant oscillations to continue despite solvent interference. The early spike potential alterations show significant proteinoid spiking flexibility that is accessible to external electrochemical manipulation.

#### 3.2.2. Modulation of Proteinoid Spiking Upon Exposure to 1.0 cm × 1.0 cm Chloroform–Embedded Filters

Exposure to chloroform for an extended period of time triggers distinct patterns of proteinoid responses, as illustrated in Figs. 7 and 26. Extended exposure to ambient accumulation for over leads to a sudden and significant reduction in inter–spike intervals. The intervals contract by more than 98%, decreasing from an average of approximately 1300 seconds to a range of 20–30 seconds (Figure 7). This hyper–excitable regime suggests crossing critical thresholds prompts greater conductivity and runaway signal cascades. Investigating the exact timing and vapour concentrations that induce hyper–excitability can reveal insights into the development of sensor materials with rapid–burst capabilities. Meanwhile, the confined spike amplitude ranges show a consistent decrease with increased chloroform insulation, measuring 0.104*±*0.011 mV upon 1×1 cm^2^ exposure (Figure 26). The distribution appears to be skewed, indicating a consistent level of proteinoid excitability despite limited intensities. The combination of acute temporal sensitization and regulated amplitude adaptation highlights the diverse proteinoid signal plasticity. By delving deeper into the textural changes and binding interactions that lead to functional modifications, we can gain a better understanding of the principles that govern vapor–induced fluctuation dynamics. Understanding the crucial tuning parameters can allow for deliberate activation of specific proteinoid states, tailored for personalised bio–electronic applications.

**Figure 7:**
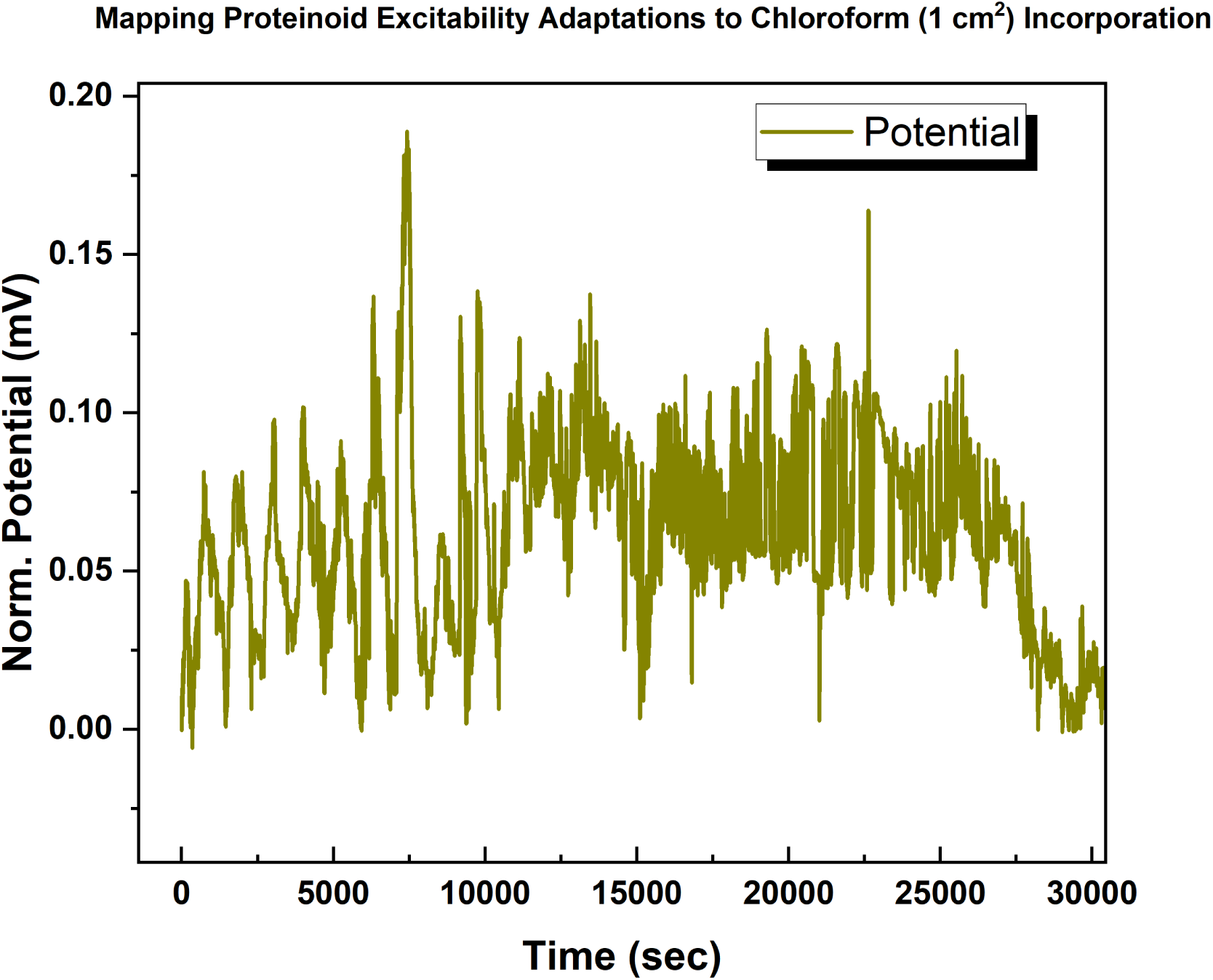
Examining the impact of extended chloroform (1*×*1 cm^2^ soaked filter paper) exposure on proteinoid spiking intervals. The graph depicts the fluctuations in bio–electric potential over a given period. At first, occasional exposure to chloroform vapour results in a reduction in the time interval between spikes, with an average of approximately 1300 seconds. Nevertheless, when exposed for an extended period of time surpassing 15000 seconds, there is a notable decrease in intervals, shrinking to a mere 20–30 seconds, which signifies a contraction of more than 98%. This sudden change implies that the combined impact of chloroform might have exceeded specific thresholds, leading to increased conductance levels and hyper–excitable dynamics. Examining the precise time intervals and levels of exposure that trigger rapid bursting could yield valuable insights into the underlying mechanisms behind intentionally inducing high–frequency patterns, potentially leading to practical applications in sensing applications.

The significant decrease in interspike intervals after prolonged exposure to chloroform in the surrounding environment is measured and presented in Fig. 21. The proteinoid spiking period was measured to be 923.00*±*107.62 seconds, indicating a significant decline of over 30% compared to the previous localised exposure (Fig. 20). The distributions display characteristics of a left–skewed Gaussian distribution with reduced kurtosis. The sudden change in spiking indicates a shift towards a highly stimulated state, with the rate of spikes increasing by more than five times. As indicated by the period metrics (Fig. 21B), the maximum intervals decrease to 1360 seconds, while the minimums approach 230 seconds between spikes. Understanding the time frame and levels of vapour concentration that trigger this heightened activity will shed light on the underlying mechanisms of this complex phenomenon. The proteinoid systems display a remarkable sensitivity to the accumulation of chloroform over time. By delving deeper into the morphologies of resultant spikes and analysing gating responses, valuable insights can be gained regarding the deliberate induction of rapid–burst exocycles, which can be highly advantageous for specific bio–sensing applications.

#### 3.2.3. Modulation of Proteinoid Spiking Upon Exposure to 3.0 cm × 3.0 cm Chloroform–Embedded Filters

Increasing the concentration of chloroform in the air on 3×3 cm^2^ filter paper causes continuing changes in proteinoid spiking properties. Positive outliers in potential (Fig. 8 and Fig. 27) may reflect these changes while preserving regular oscillatory patterns (Fig. 22). Spike potentials show a significant decrease, with average values of 0.091*±*0.008 mV, suggesting a reduction of nearly 90% when compared to the control. The potential distribution shifted to the right indicates a significant change in conductivity, however there are brief exaggerations that exceed the maximum values of the control. Under these conditions, the time intervals between spikes likewise decrease (Fig. 22), with average values of around 1000 seconds. The fact that there is a continuous pattern even at the lowest duration of approximately 550 seconds and the longest duration of approximately 1500 seconds, on the other hand, illustrates the durable nature of these artificial stimulation systems in the presence of vapour phase disturbances. Analysing the morphological changes in waveform adaptations observed by statistical profiling would provide a better understanding of the tuning principles that govern long–term stimulus responsiveness.

**Figure 8:**
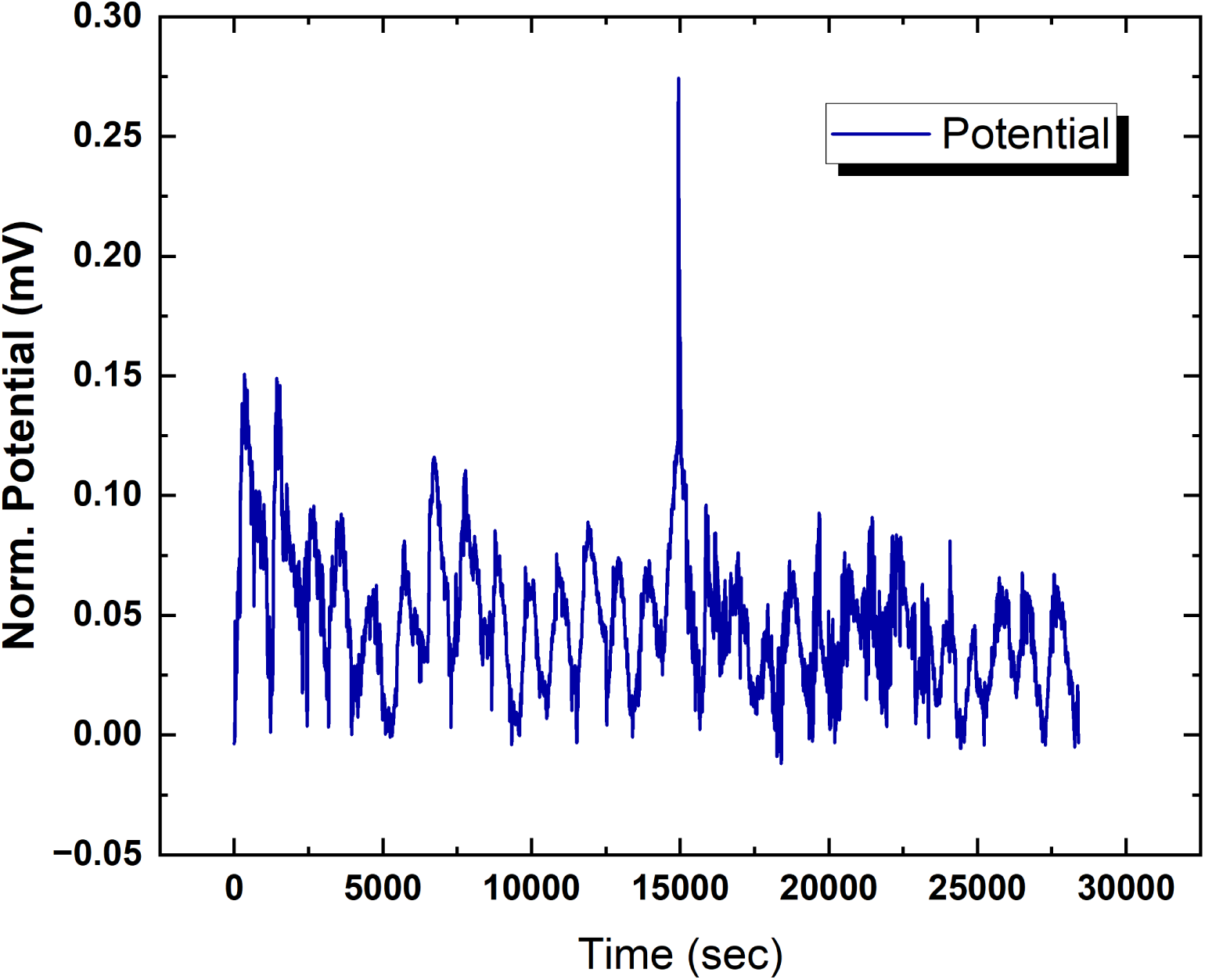
Bio–electric potential dynamics with increased exposure to a 3*×*3 cm^2^ chloroform–soaked filter. The plot shows a significant reduction in the mean spike amplitude of 0.091*±*0.008 mV (minimum 0.049 mV; maximum 0.274 mV) across a 15–minute period of modulated spiking oscillations in the plot. Positive skewness (2.99) and high kurtosis (12.93) in the data distribution point to an outlier–pronounced distribution at higher doses.

### 3.3. Solvated Chloroform Response Dynamics

Exposing thermal proteinoid polymers directly to chloroform at a concentration of 25mg/L caused significant but controlled changes in the patterns of emerging excitation waves (Fig. 9–28). Average spike amplitude detected 0.065*±*0.009 mV - demonstrating 90% attenuation relative to baseline (Fig. 28). Simultaneously, the frequency became highly unpredictable with occasional contraction, reaching temporary intervals as short as 69.80 seconds (Fig. 23). Despite the disturbance, there are consistent and rhythmic fluctuations that demonstrate the impressive ability of proteinoids to recover and adapt (Fig. 9). These fluctuations are presumably a result of self–adaptation mechanisms that actively maintain conductive structures. Elucidating the precise order of binding interactions, structural changes, and controlled shifts in conductance that contribute to the observed functional persistence may enhance the development of adaptable bio–electronics materials based on biological principles. Gradient titrations can also uncover key concentration thresholds that trigger either uncontrolled collapse or preserved excitability. Utilising limited but viable ranges creates possibilities for deliberate and adjustable destabilisation, which in turn activates quick and intense sensory capabilities that are helpful in identifying specific organic substances. Ultimately, the demonstrated adaptive plasticity of proteinoids provides fundamental templates for developing computer circuits that are artificially robust and responsive to the environment.

**Figure 9:**
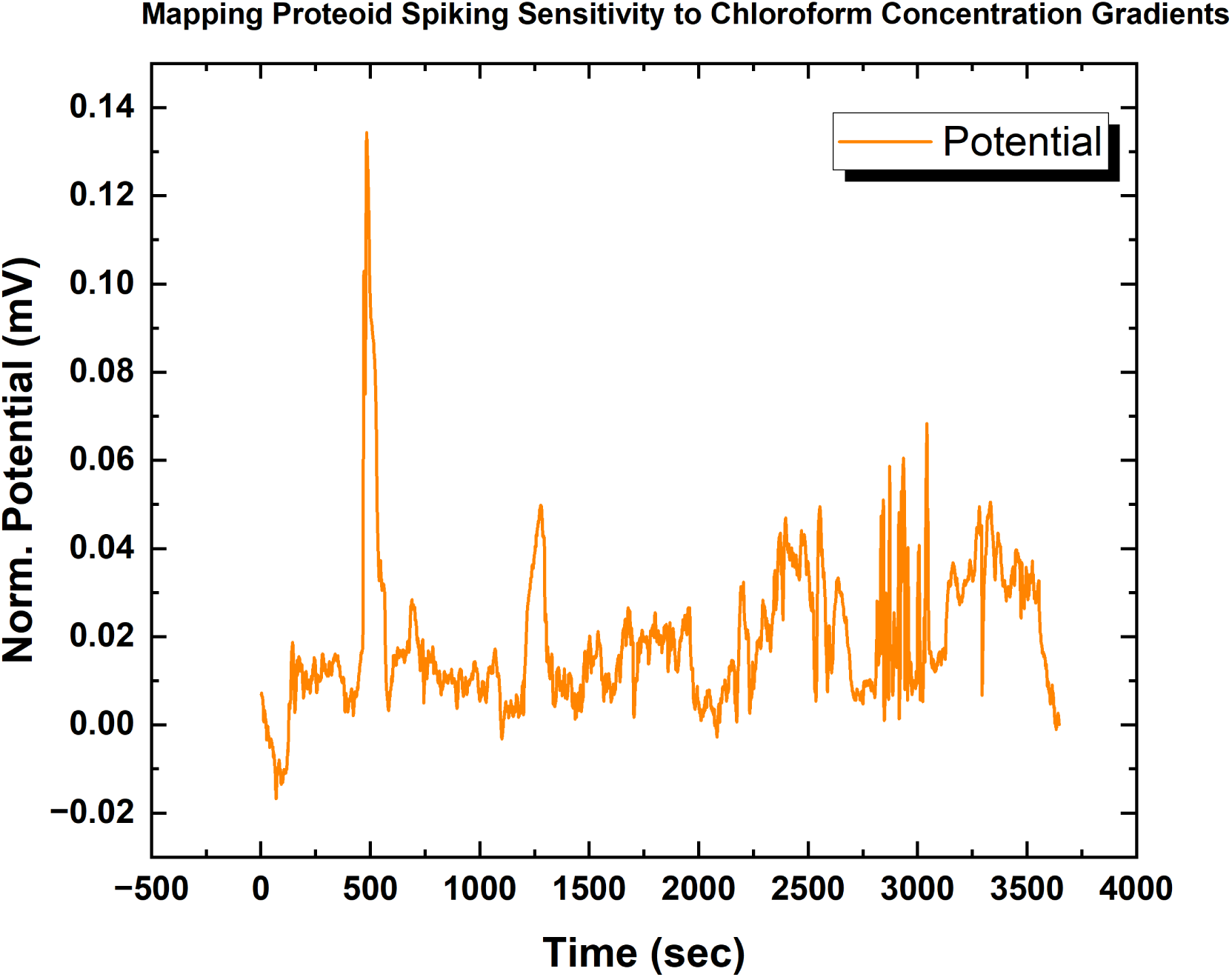
Spiking activity of proteinoids diluted with 25mg/L chloroform. The recorded waveform displays changes in spiking patterns over a time period of 900 seconds, characterized by a notable decrease in mean amplitude of 0.065*±*0.009 mV compared to control conditions. Variability in periodicity is also evident, with intervals ranging from 69.80 to 623.00 seconds. Interestingly, the observed oscillations persist despite the disruption, indicating the operation of inherent homeostatic mechanisms that maintain functional conduction pathways.

**Figure 10:**
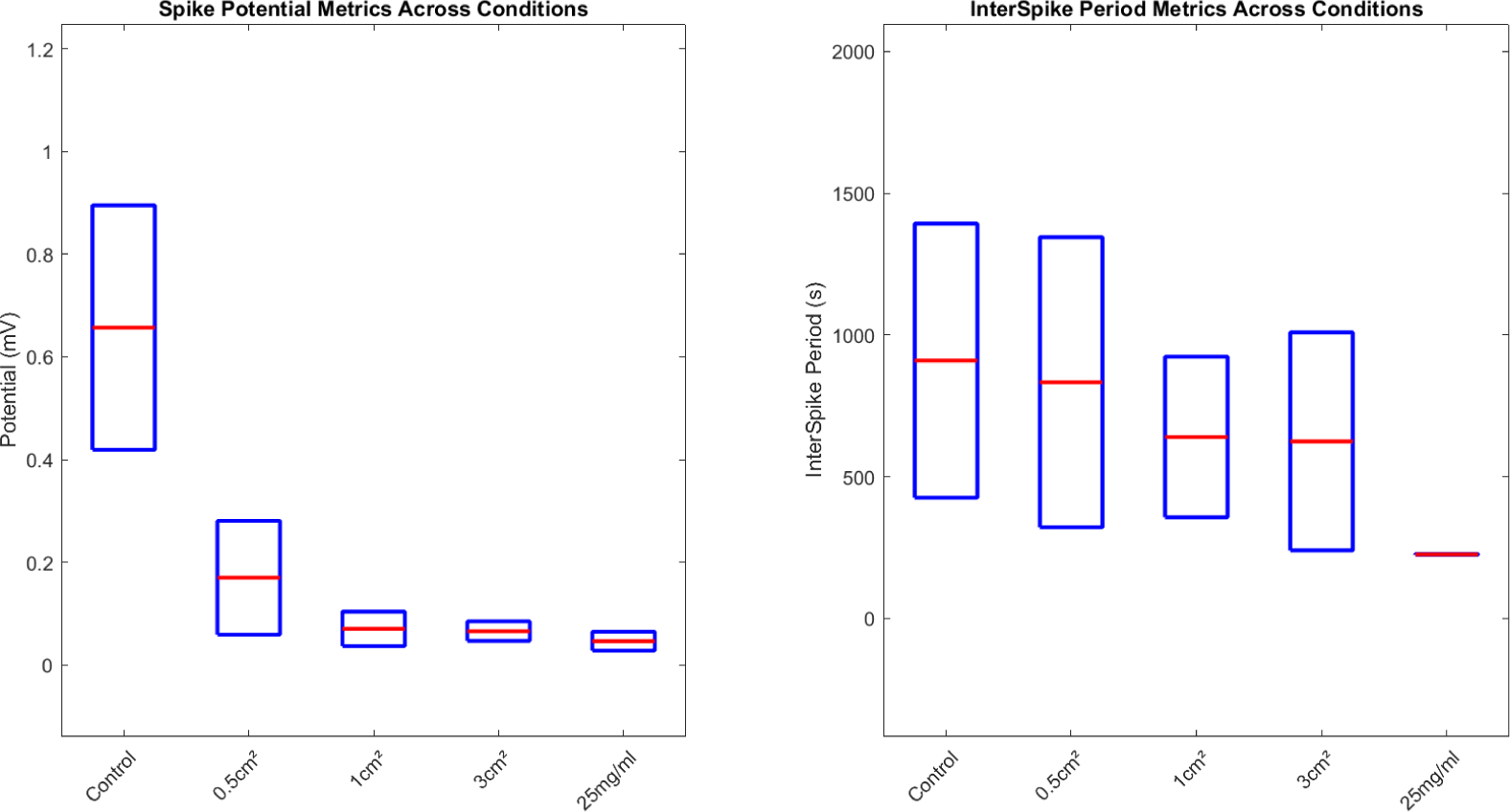
Interspike period measurements and spike potential metrics in proteinoids depicted across different chloroform exposure levels. Under control conditions and chloroform concentrations of 0.5 cm^2^, 1cm^2^, 3cm^2^, and 25mg/mL, each box plot represents the distribution of interspike intervals (in seconds) and the distribution of spike potentials (in millivolts). Each box delineates the IQR, with the middle line denoting the median value, demonstrating the data’s distribution and variability. The broad spread in the ’Control’ and ’1cm^2^’ conditions signify a wider variation in inter–spike period measurements.

**Figure 11:**
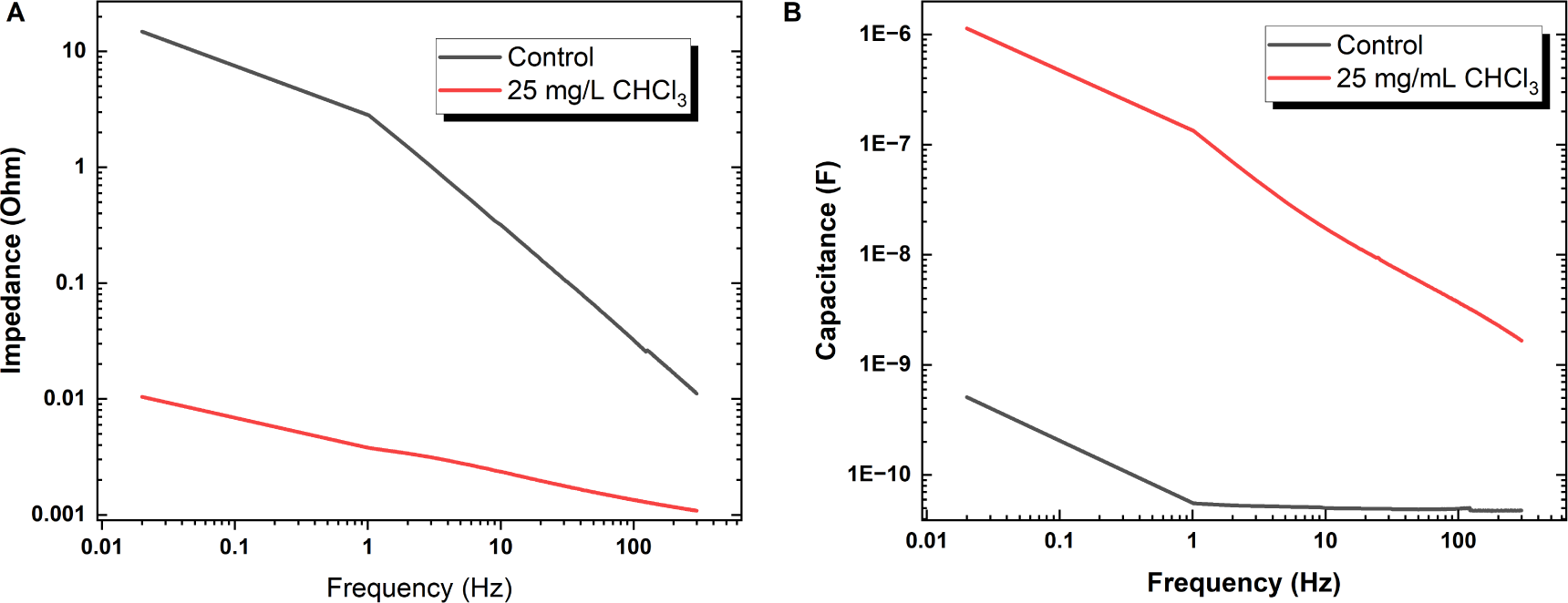
Frequency response of electrical characteristics in control and chloroform–exposed proteinoid samples. (A) Capacitance exhibits a logarithmic declining trend with increased frequency for both control (mean = 0.050*±*0.027nF) and 25mg/L chloroform exposed (0.087*±*0.66nF) cases. (B) Similarly, impedance progresses logarithmically for control (0.115*±*0.875 Ω) and exposed (1.3956*×* 10*^−^*^3^ *±* 6.4324*×*10*^−^*^4^ Ω) conditions. In both metrics, chloroform introduction sharpens dynamic range and reduces the regularity of impedance spikes observed inherently, indicating regulated energy storage and transfer capacities. Recording dose–responsive trait alterations as frequency spectra shift could direct intentional tuning of electrical properties for specialized bio–electronic applications.

**Figure 12:**
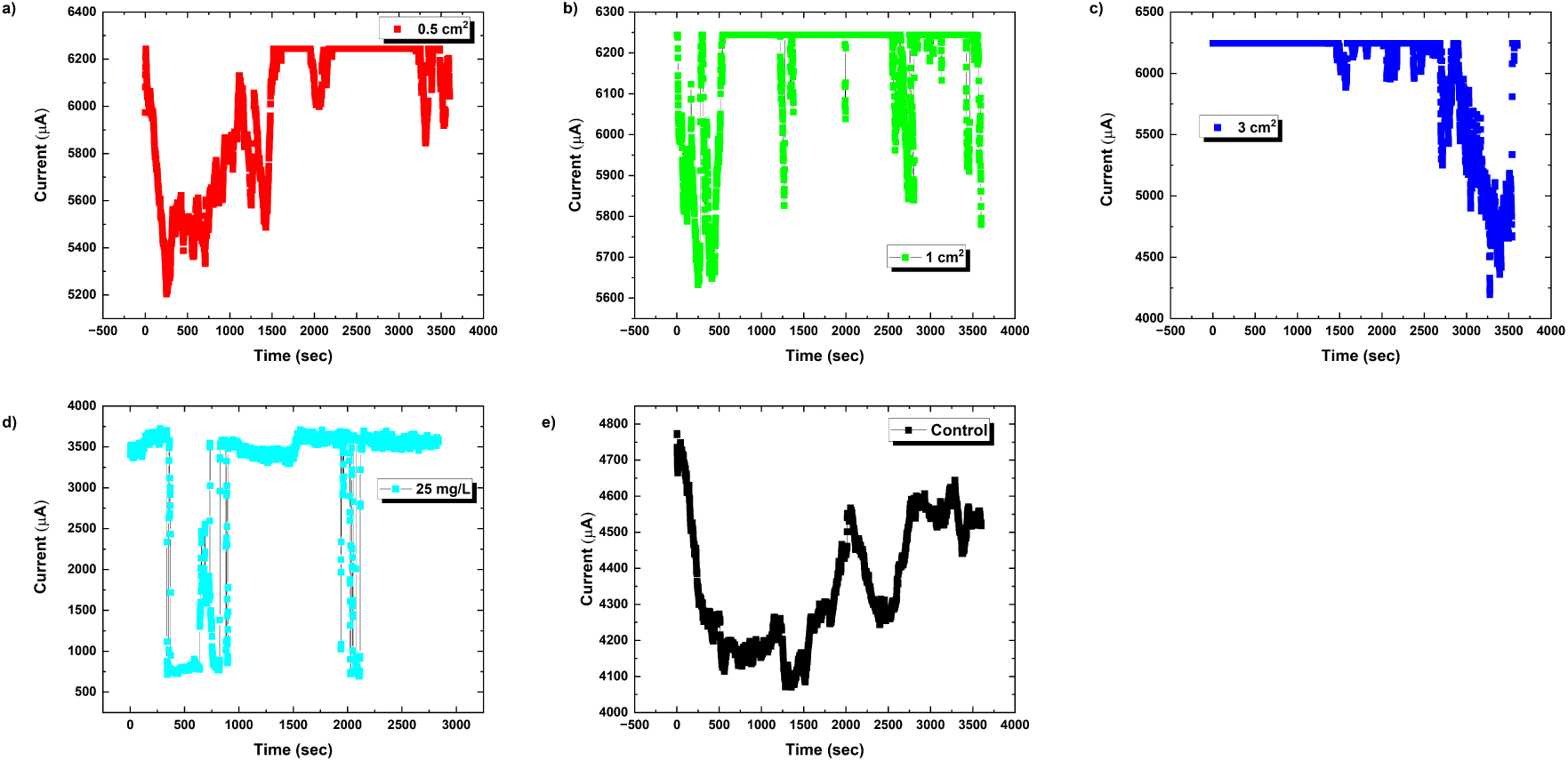
The chronoamperometric current responses of L–glutamic acid:L–arginine (L-Glu:L-Arg) proteinoids were measured over time as the chloroform (CHCl_3_) concentrations increased. A control was included for comparison. The equilibrium duration is 0 seconds, the applied DC voltage is 0.1 volts, and the sampling interval is 1 second. Proteinoids exhibit temporary increases in conductivity that are influenced by changes in solvent levels, resulting from alterations in conformation and morphology. The rapid relaxation of memristive dynamics within subsecond timeframes suggests the stability of peptide microspheres.

**Figure 13:**
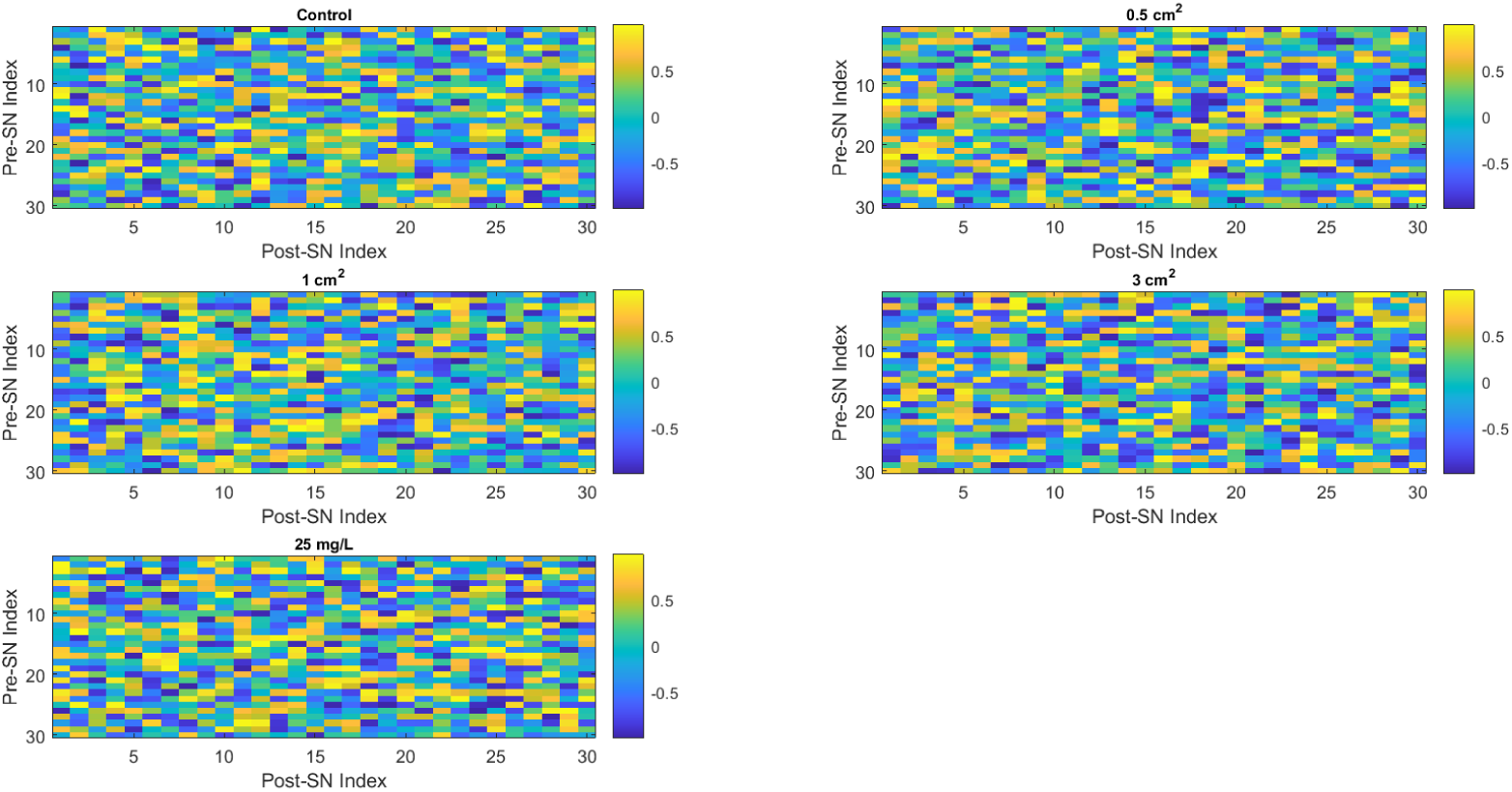
Heat maps of initial synaptic weights for various conditions. Each of the five subplots depicts the initial synaptic weights for each condition: control, 0.5 cm^2^, 1 cm^2^, 3 cm^2^, and 25 mg/L chloroform exposure. The synaptic weights are generated at random and displayed as a colour–coded matrix, with the colour intensity corresponding to the synaptic weight value. The *x* and *y* axes show post–synaptic and pre–synaptic neuron indices, respectively. These heat maps provide a preliminary overview of neural activity for every scenario.

**Figure 14:**
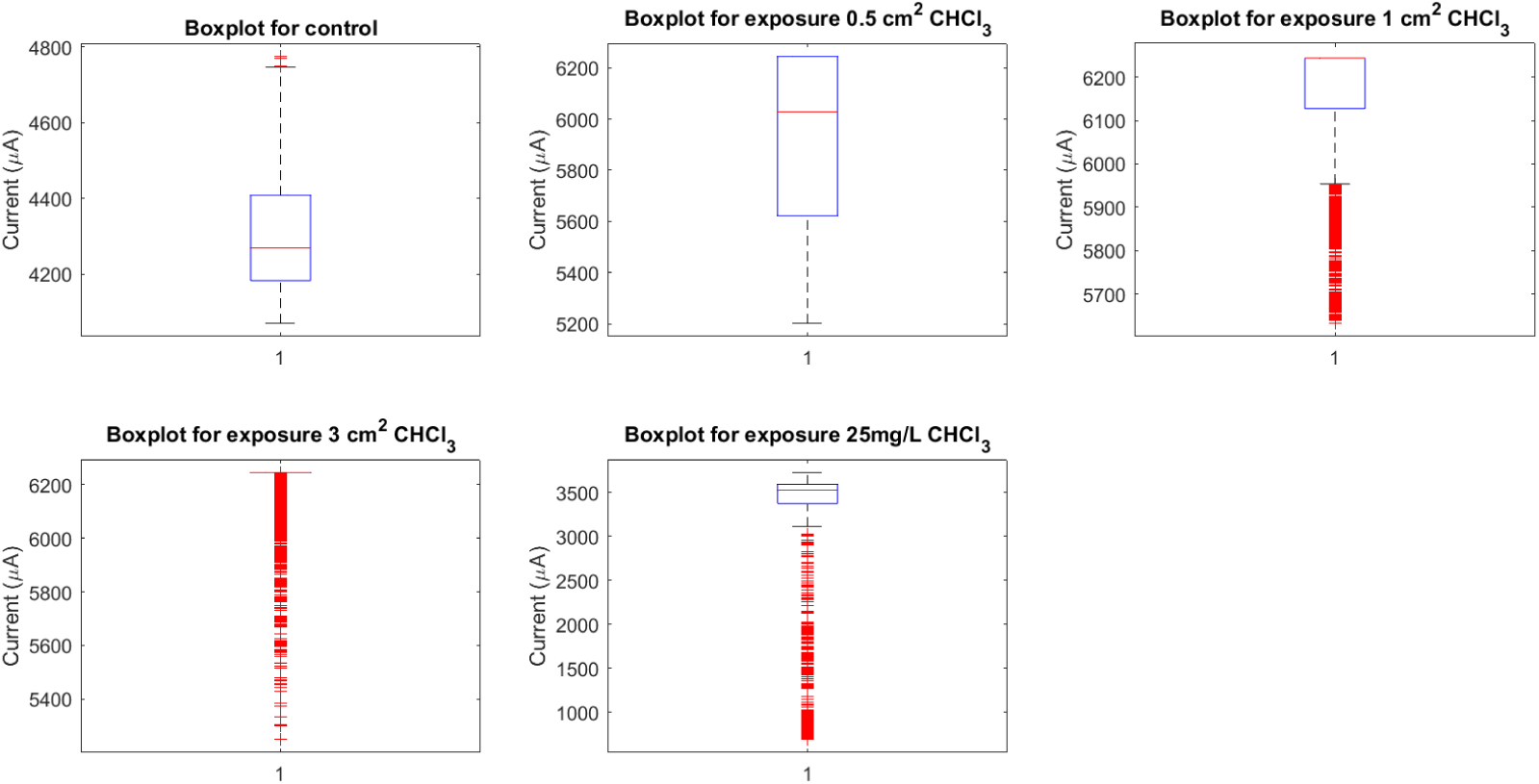
Current distributions under different conditions. The image depicts boxplots of the currents for five conditions: control, 0.5 cm^2^, 1 cm^2^, 3 cm^2^, and 25 mg/L chloroform exposure of proteinoids. Each box’s central line represents the median current, while the top and bottom bounds represent the upper (Q3) and lower (Q1) quartiles, respectively. The whiskers extend to the most extreme data points that are not deemed outliers, which are represented individually as dots. The notches around the median represent the median’s uncertainty. Mean currents (A) for control, 0.5 cm^2^, 1 cm^2^, 3 cm^2^, and 25 mg/L are around 4305, 5931, 6157, 6202, and 3046 *µ*A, respectively, indicating a considerable drop in the current following chloroform exposure at 25mg/L.

**Figure 15:**
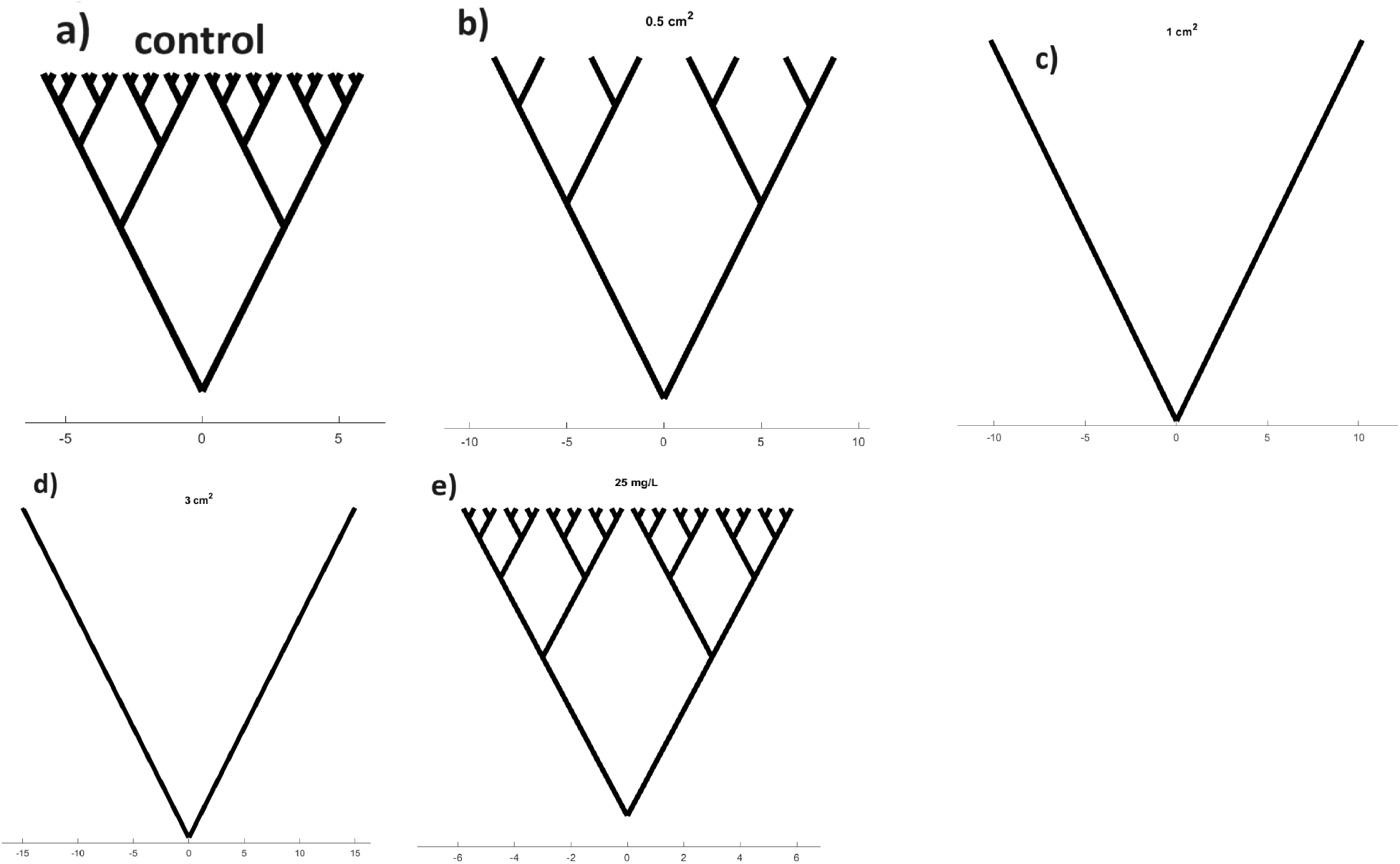
Binary tree pathways conceptually describe changes in the complexity of the interpreted proteinoid network following chloroform exposure. The subfigures depict the following conditions: a) Control, b) 0.5 cm^2^, c) 1 cm^2^, d) 3 cm^2^, and e) 25 mg/L. As the concentration of chloroform increases, trees undergo a transition from initially having more complex branching patterns (a) to simpler forms with fewer forks (b,d). This suggests that there is an emergence of enhanced structural order in proteinoid assemblies. However, when exposed to a concentration of 25 mg/L, the branches undergo recomplexification, indicating a possible degradation of organised dynamics under extreme conditions. Reduced maze–like trees, albeit abstract, symbolise the observed increase in order resulting from solvent interactions, which could perhaps be attributed to the molecular initiation of self–construction. The variations in complexity observed at different levels of exposure emphasise the ability to adjust proteinoid computational structures through external chemical regulation.

**Figure 16:**
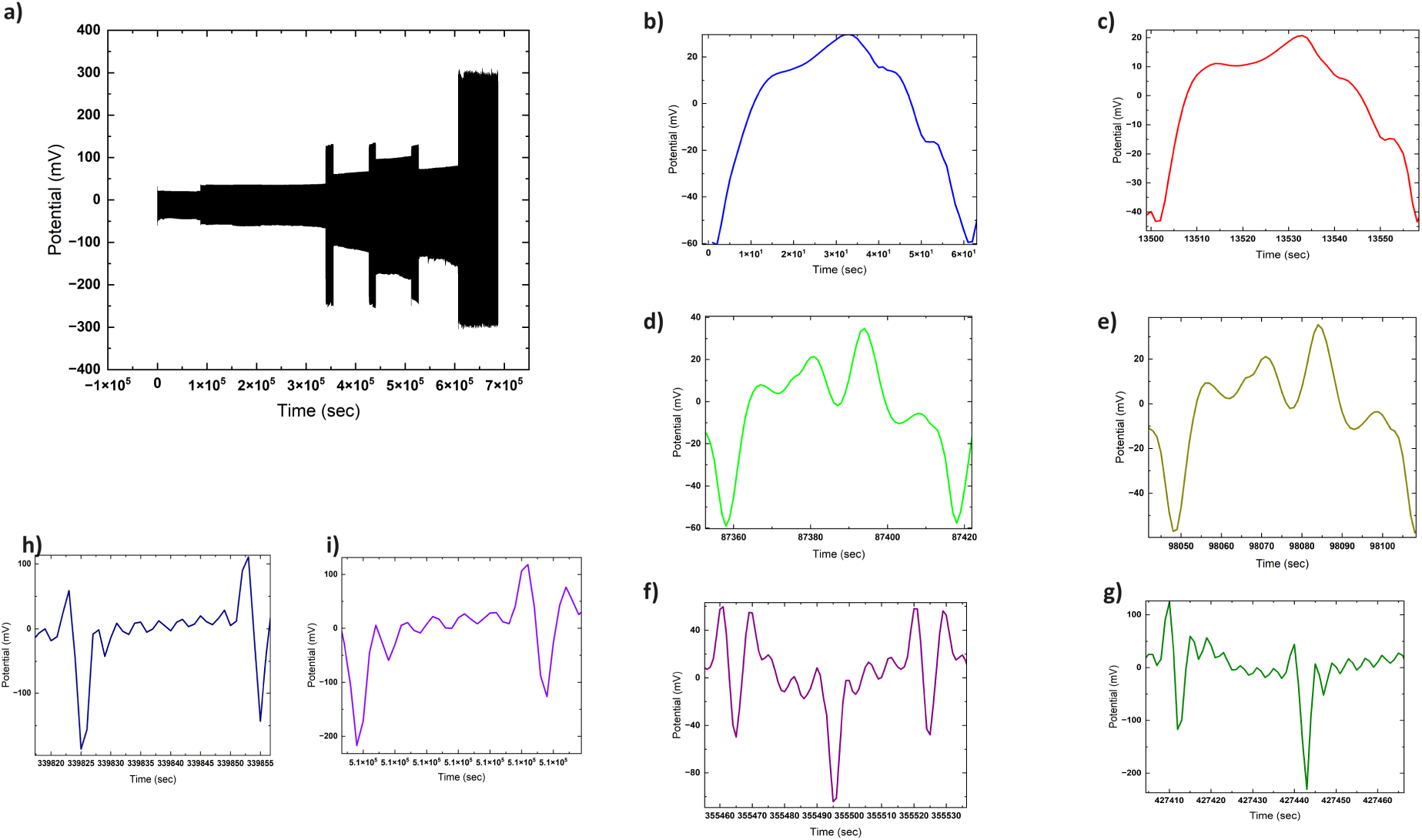
Sample recordings of output voltage signals generated by a 25mg/L chloroform–modulated proteinoid assembly acting as a voltage–stimulated proto–brain system. Trace (a) shows the overlay of complex output signal over time containing multiple embedded frequency components. Enlargements highlight specific output signal harmonics parsed from the composite response: (b) 1st (fundamental) harmonic, (c) 2nd harmonic, (d) 4th harmonic, (e) 5th harmonic, (f) 8th harmonic, (g) 14th harmonic, (h) 16th harmonic, and i) 13th harmonic. Presence of higher–order mapped harmonics in the proteinoid voltage output demonstrates nonlinear transformational responses analogous to neural–system kernels. Sensitive input tuning may enable selective filtering and computational pathway–gating by harmonically complex bio–derived signal processing elements.

**Figure 17:**
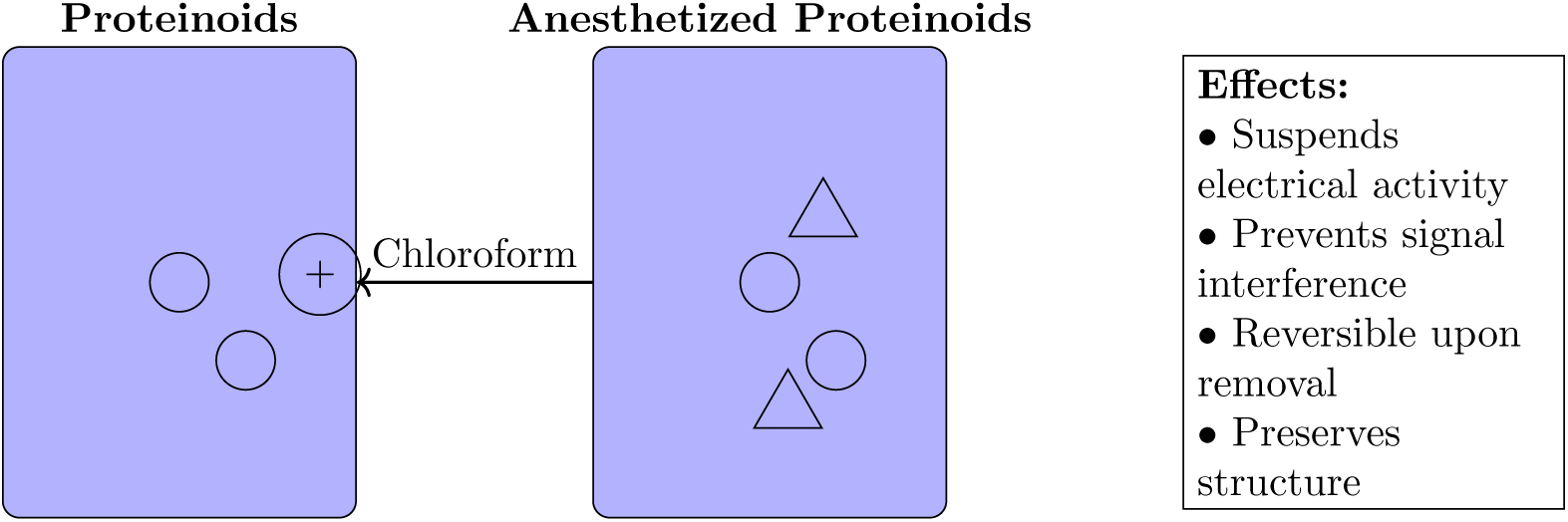
Anesthesia of proteinoids using chloroform.

### 3.4. Concentration–Dependent Effects

Tables 1 and 2 summarise important measured proteinoid spiking potential and periodicity properties throughout steadily increasing chloroform exposures encompassing vapour phase and direct solvation introduction.

**Table 1:**
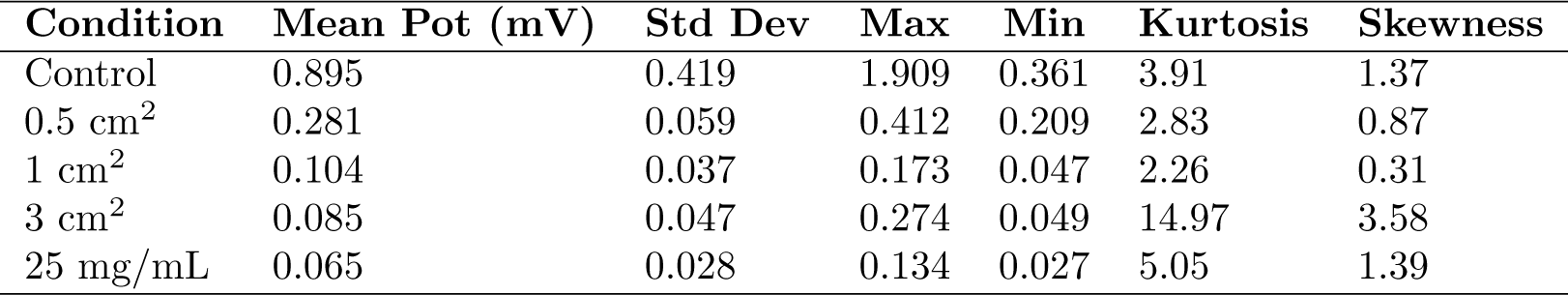
Spike potential metrics across conditions.

| Condition | Mean Pot (mV) | Std Dev | Max | Min | Kurtosis | Skewness |
| --- | --- | --- | --- | --- | --- | --- |
| Control | 0.895 | 0.419 | 1.909 | 0.361 | 3.91 | 1.37 |
| 0.5 cm<sup>2</sup> | 0.281 | 0.059 | 0.412 | 0.209 | 2.83 | 0.87 |
| 1 cm<sup>2</sup> | 0.104 | 0.037 | 0.173 | 0.047 | 2.26 | 0.31 |
| 3 cm<sup>2</sup> | 0.085 | 0.047 | 0.274 | 0.049 | 14.97 | 3.58 |
| 25 mg/mL | 0.065 | 0.028 | 0.134 | 0.027 | 5.05 | 1.39 |

As demonstrated in Tab, 1, average spike intensity gradually decreases with increasing chloroform concentration, with mean values decreasing from 0.895 mV to 0.065 mV from control to 25 mg/mL conditions, indicating almost 90% damping.

Simultaneously, amplitude distribution widths narrow significantly, restricted to smaller ranges between peak and minimal spikes. Similarly, increased chloroform levels cause a shortening of interspike intervals (Tab. 2), reflecting increased excitation states. In essence, spikes compression in conjunction with rhythmic coordination highlights a suppression of bio–electric disorder in the face of increasing solvent perturbation.

**Table 2:** Interspike period metrics across conditions.

| Condition | Mean Period (s) | Std Dev | Max | Min | Kurtosis | Skewness |
| --- | --- | --- | --- | --- | --- | --- |
| Control | 1392.86 | 425.83 | 1952.38 | 595.24 | 1.95 | -0.36 |
| 0.5 cm<sup>2</sup> | 1344.85 | 321.53 | 1871 | 882 | 1.82 | 0.41 |
| 1 cm<sup>2</sup> | 923 | 356.92 | 1360 | 230 | 2.67 | -0.87 |
| 3 cm<sup>2</sup> | 1008.75 | 240.08 | 1505 | 718 | 3.46 | 0.95 |
| 25 mg/mL | 228.2 | 223.91 | 623 | 69.8 | 2.08 | 0.92 |

When comparing the control and 25 mg/L groups, the periodicity kinetics increase by more than 80%. Sharper rhythmic coordination emerges in conjunction with compressed waveform volatility, indicating more systematic ion conductance underneath the waveforms. Nonetheless, sustained spiking demonstrates persisting proteinoid bio–electricity with non–Gaussian statistical features, indicating a retained nonlinear dynamics despite dose escalation up to solvent saturation. Following morphological changes at chloroform junctions can now reveal particular mechanisms that support regulated waveform adaptation.

As shown in Fig. 10, boxplots summarize the distribution of key spiking metrics across incrementally elevated chloroform regimes. For inter–spike periods, control proteinoids exhibit the broadest spread spanning 595.24 to 1952.38 seconds. This likely reflects intrinsic bio–electric variability in the absence of external disruption. Upon initial localised vapour exposure (0.5 cm^2^), marginal interval tightening appears, progressively contracting further under more extensive coverings (1 cm^2^ and 3 cm^2^). Finally, direct solvent contact intensely heightens rhythmic rates, driving mean periods down to 228.2 *±* 223.91 seconds.

Similarly for amplitudes, higher intensity distributions prevail in control samples up to 0.9 mV, sequentially declining under rising chloroform presence down to 0.065*±*0.028 mV. Tighter, lower intensity profiles emerge denoting constrained excitation and pathway conduction.

Figure 10 thus quantifies chloroform’s dose-dependent impacts in concentrating emergent rhythms yet attenuating signal output – pointing to re–optimisation not collapse. Further interrogation of compensatory mechanisms sustaining modulated waveforms could direct fabrication of resilient bio–electronic materials.

#### 3.4.1. Tuning Resistance and Reactance in Proteinoid Assemblies Through Chloroform Incorporation

Figure 11, sweeping proteinoid sample exposure to varied input signal frequencies reveals key electrical traits altered upon chloroform incorporation. Most prominently, a logarithmic decline in dynamic capacitance manifests in both control and exposed samples, with mean values of 0.05 nF and 0.087 nF respectively. The heightened variability and range compression with chloroform suggests regulated rather than disrupted energy storage modalities. Similarly, impedance spectra progress logarithmically from 0.115*±*0.875 Ω in control state down to 1.3956× 10*^−^*^3^ *±* 6.4324×10*^−^*^4^ Ω when exposed. In effect, the programmed proteinoid systems exhibit adaptive self–correction of electrical characters rather than waveform collapse when confronted with chloroform disturbances. Further elucidation of the precise binding mechanisms and morphological adaptations eliciting enduring, regulated bio–electric identities could direct fabrication of resilient, specialized conductive materials.

#### 3.4.2. Concentration–Dependent Modulation of Proteinoids–Chloroform Memristive Dynamics Characterized by Chronoamperometry

Chronoamperometry reveals brief electrical responses of proteinoids systems, as illustrated in Figure 12 for L–glutamic acid:L–arginine under increasing chloroform concentration. The memristive spike and recovery dynamics show that solvent interactions cause transitory conductivity enhancement.

Figure 12 depicts the time–dependent current for various situations, with time on the *x*-axis and current on the *y*-axis. The present responses indicate various patterns, indicating the effect of proteinoid exposure to chloroform under varied settings.

Let us analysis the experimental results in the framework of imitated neural networks. Fig. 13 shows an analysis of the neural networks’ initial synaptic weights. The heat maps of generated synaptic weight matrices operate as a symbolic representation of the patterns of neural connections that could arise in unconventional computing systems based on proteinoids. These 2D visualisations illustrate potential variations in neuronal organisation under different chemical conditions by representing proteinoids suspensions as a concrete neuromorphic substrate, albeit the notion is still in its early stages. For example, the noticeable variations in colour intensity and gradient distributions may indicate the formation of different network structures and collective behaviours. We expect to observe more clustering in control situations, while chloroform exposure is likely to result in higher randomness. Continuing simulations that incorporate these patterns of connection and function into models of spiking neural networks can assist in forecasting the emerging computing abilities. Furthermore, by establishing a correlation between the characteristics of the heat map and the observed conductivity behaviours and recovery durations in experiments (as shown in Fig. 13), we can get insights into the relationship between short-term and long-term plasticity manifestations and the development of simulated architecture. In addition to static visualisation, the inclusion of temporal changes towards stable end states can enhance our understanding of how proteinoid systems maintain a balance between order, chaos, and adaptable bio-derived computing.

The Eq. 1 can expresses the current transition as follows:

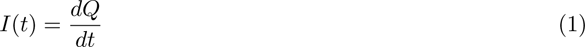

where *I*(*t*) signifies the current and *Q* symbolises charge. One can calculate the current by differentiating the charge with regard to time. The shifting current patterns under varied settings (shown in Fig. 12) indicate a change in charge flow rate due to variable levels of chloroform exposure.

The synaptic weight matrix (*W*) can be first expressed in neural network modelling as:

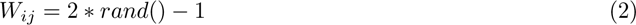

where (*W_ij_*) represents the synaptic weight from the (i)th presynaptic neuron to the (j)th postsynaptic neuron and rand() returns a random value between –1 and 1. As illustrated in Fig. 13, the effect of chloroform exposure causes a clear difference in the initialisation of synaptic weights.

The simulation of a neural network was implemented through generating artificial neurons and connecting them with “synaptic” weights. For each condition analysed, we created a network including 30 simulated neurons. The neural activity was correlated with a binary temporal coding, which was obtained from the time–dependent current signals seen in our investigations. In order to create these temporal codes, a sampling threshold of 1e-4 *µ*A was established. Any recorded current signal above this threshold was considered as neural activity and designated as a state of ’1’. Subsequently, we randomly chose a subset of these data points, making sure that the size of the group was inversely related to the number of artificial neurons. This approach added a realistic aspect to our model by incorporating the intrinsic unpredictability found in biological systems. We assigned a set of initial synaptic weights to each neuron in the network. These weights were determined arbitrarily as random values within a range of –1 to 1. The weights were specifically intended to adapt themselves iteratively according to a predetermined learning rule, allowing the network to learn progressively over time. The initial allocation of these synaptic weights was shown as a heatmap for each condition, providing us with a visual understanding of the starting state of the network prior to any learning taking place.

Finally, the chloroform appears to have a distinct influence on the electrical properties of the proteinoid, which is detectable through changes in current and seen in the variation of synaptic weights in our model’s neural network.

Neuronal currents exhibit varied distribution characteristics under the five different situations, as shown in Fig. 3.4.2. Notably, the mean current increases incrementally from the control condition to the 3 cm^2^ chloroform exposure condition. However, when proteinoids are exposed to chloroform at a concentration of 25 mg/L, the mean current drops significantly to around 3046 *µ*A. The boxplot’s quartiles and whiskers provide a detailed view of the data dispersion and probable outliers for each scenario. The median neuronal current has a similar pattern to the mean current. The significant decrease in median current upon chloroform exposure demonstrates the significant impact of chloroform on the electrical characteristics of the proteinoid.

Lempel–Ziv (LZ) complexity is a measure that quantifies the level of randomness in finite sequences or patterns [61]. As observed in the synaptic weight maps (Fig 13), the LZ complexity declined as the chloroform exposures increased up to 3 cm^2^, and then showed a minor recovery at 25 mg/L. The control instance demonstrated the highest LZ score of 4.7399, suggesting a pattern that is almost maximally complex. A decrease in LZ complexity indicates an increase in the organisation and predictability of the created connection graphs. This indicates that when solvent concentrations increase, the architecture of neurons tends to shift from disorder to a more organised and localised clustering. Nevertheless, once a specific limit is surpassed, excessive disruption of proteinoids can lead to the degradation of organised assembly. In general, the fluctuations in non–monotonic LZ complexity are associated with the possibility of disturbance followed by the restoration of self–organised circuit development when influenced by chloroform. The quantitative complexity metrics enhance the analysis of induced folding modifications and their organisational propagation throughout architectural development, providing better insights than visual map inspection alone. Table 3 summarises the Lempel–Ziv complexity results.

**Table 3:** Lempel–Ziv complexity under different chloroform exposure conditions.

| Condition | LZ Complexity |
| --- | --- |
| Control | 4.739922 |
| 0.5 cm<sup>2</sup> | 2.790569 |
| 1 cm<sup>2</sup> | 1.383151 |
| 3 cm<sup>2</sup> | 1.002987 |
| 25 mg/L | 4.578150 |

The image depicts boxplots of the currents for five conditions: control, 0.5 cm^2^, 1 cm^2^, 3 cm^2^, and 25 mg/L chloroform exposure of proteinoids. Each box’s central line represents the median current, while the top and bottom bounds represent the upper (Q3) and lower (Q1) quartiles, respectively. The whiskers extend to the most extreme data points that are not deemed outliers, which are represented individually as dots. The notches around the median represent the median’s uncertainty. Mean currents (A) for control, 0.5 cm^2^, 1 cm^2^, 3 cm^2^, and 25 mg/L are around 4305, 5931, 6157, 6202, and 3046 *µ*A, respectively, indicating a considerable drop in the current following chloroform exposure at 25mg/L.

The image depicts boxplots of the currents for five conditions: control, 0.5 cm^2^, 1 cm^2^, 3 cm^2^, and 25 mg/L chloroform exposure of proteinoids. Each box’s central line represents the median current, while the top and bottom bounds represent the upper (Q3) and lower (Q1) quartiles, respectively. The whiskers extend to the most extreme data points that are not deemed outliers, which are represented individually as dots. The notches around the median represent the median’s uncertainty. Mean currents (A) for control, 0.5 cm^2^, 1 cm^2^, 3 cm^2^, and 25 mg/L are around 4305, 5931, 6157, 6202, and 3046 *µ*A, respectively, indicating a considerable drop in the current following chloroform exposure at 25mg/L.

The schematic (Fig. 15) is designed to visually represent the overall trend of increased order that emerges in the self–organised dynamics of proteinoids as the chloroform levels increase. The purpose behind generating the binary tree abstraction, as depicted in Fig. 15, lies in its ability to cognitively embody the nonlinear association between the morphological complexity of adjustable proteinoids and the observed concentration levels of chloroform exposure during experimentation. To be specific, an initially careful and controlled addition of a minuscule amount of the solvent contributes significantly to the formation of a highly consistent and orderly structure throughout the process of molecular self–assembly. This is reflected in a marked decrease in the branching complexity. Notwithstanding, this process of simplification begins to retrogressively reverse at higher saturation levels, suggesting that supersaturation adversely impacts the organised dynamics of the system. We thereby put forth the proposition that proteinoid systems exhibit a labyrinthine configuration optimisation landscape, influenced by interactions that do not necessarily adhere to a linear pattern. Furthermore, the visualisation, while qualitative, inspires quantitative investigations into precisely defined solution parameters for optimally hatching desired bio–architectures – whether dendrimers, fractals, or other uniform meshes ideal for unconventional computing.

Nevertheless, the particular intricacy of branching patterns represents one way to quantify the randomness of patterns, but there are other valid interpretations of the model as well. We believe that the importance of this lies in demonstrating the possibility of using chemical parameter alterations alone to guide proteinoid structural bifurcations towards desired computational geometries. By enhancing solution control and gaining a deeper understanding of how macroscale assembly is influenced by molecular triggers, it is possible to create proteinoid architectures that closely resemble basic living cells.

### 3.5. Elucidating Input–Output Relationships of Chloroform–Tuned Proteinoid Network Bio–interfaces Under Periodic Voltage Harmonic Forcing

Complex signals and harmonic analysis are tools for quantifying nonlinear system behaviours. Any periodic waveform can be represented as a sum of distinct harmonic frequency components:

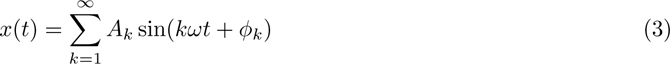

where *A_k_*and *ϕ_k_* are harmonic amplitude and phase, *ω* is base frequency, and *k* indexes each harmonic component.

Figure 16 illustrates that solvent–modulated proteinoid systems produce a diverse range of harmonic outputs when exposed to different voltage waveforms. The emerging higher–order harmonics exhibit nonlinear transformational reactions that are similar to neural circuits [62, 63]. Tracing the source of particular harmonics allows us to gain understanding of the corresponding molecular processes occurring in their natural environment — such as ion movement, membrane polarisation, conformational changes, or structural rearrangements. In the end, utilising input harmonics that efficiently drive certain proteinoid harmonic outputs could potentially allow for precise computational filtering and pathway–gating in bio–derived unconventional computing.

The rich mosaic of harmonic content embedded within the observed proteinoid response signals, as depicted in Fig. 16, shows that similarly elaborate processing is possible. Isolating the origins of unique harmonic signatures remains a challenge. However, the diversity implies that protonic, electronic, and conformational movements all interact and contribute in different ways. Excitingly, using multifrequency harmonic forcing as a computational tuning parameter may soon allow for the manipulation of desirable collective dynamics.

## 4. Potential molecular mechanisms

Chloroform has been extensively used as an inhalation anaesthetic. It causes organisms to lose consciousness by entering and affecting the lipid bilayer membranes, thereby disrupting important signalling pathways [64]. Studies have specifically demonstrated that chloroform interferes with specialised lipid raft microdomains that are crucial for the localization of enzyme activity and cell communication mechanisms [65, 66, 67]. The indiscriminate buildup of chloroform in membranes hinders the proper functioning of enzymes such as phospholipase D2 (PLD2), which regulate brain signalling molecules. Moreover, the focusing of substances at the membranes of nerve cells disrupts ion channels such as TREK–1, which necessitate accurate control in order to transmit electrical signals that are fundamental to cognitive function and alertness. By utilising these established methods of membrane disruption, chloroform and similar anaesthetic chemicals can effectively reduce consciousness and brain activity that are crucial for awareness and cognitive functions in living organisms.

Positioning it in stark contrast to living cell membranes, where chloroform exposure results in the disruption of ion channels and the eventual electrical isolation of the bilayer, the proteinoid microspheres tested within this study have been observed to maintain voltage oscillations, despite being subjected to similar levels of solvent exposure. This striking incongruity suggests that there exist fundamental differences between the mechanisms of excitation of the two involved systems. The persisting harmonics displayed by the proteinoids hint at the dominance of bulk effects arising from various factors - the pressure strains brought upon by swelling, the formation of temporary pores, and alterations in the dynamics of molecular clustering - over the inhibition of ion movement. In a proteinoid system, the small soluble chloroform molecules may penetrate the peptide matrix, which has been thermally cross-linked, in their attempts to fragment it. This action results in the oscillations in the proto–cell wall, causing it to alternately contract and expand.

How can chloroform cause proteinoid microspheres to conduct and oscillate, unlike lipid bilayer cell membranes, which are insulating?

As a preliminary attempt to generate a hypothesis, we proffer potential explanations which focus on the permeation of the proteinoid matrix, the phenomena associated with swelling, the disruption of interior dynamics, along with the absence of surface ion channel analogues.

Potential Mechanisms:

- **Transient pore formation** Small chloroform molecules infiltrate and cause temporary defects in cross–linked peptide matrix.
- **Swelling strain effects** Proteinoid microsphere’ wall dimensions increase or decrease with solvent attack.
- **Modulated molecular clustering** Disrupts interior packing dynamics and charge mobility pathways.
- **Lack of ion channel disruption** Unlike cells, no insulating from impedance of selective ion flow.
- **Bulk matrix effects rather than surface phenomena** Deep permeation into peptide structure underpins observed dynamics.

This in–depth understanding of the modus operandi of proteinoid microspheres can be vitally instrumental in deciphering their versatility and potential utilities in the realm of unconventional computing. It propels the investigation a step forward towards the efficient utilisation and manipulation of these bio–architectures in the development and designing of innovative computational systems.

## 5. Discussion

The results of our research provide compelling evidence that increasing chloroform levels progressively alter the electrical signalling of proteinoids. We observed the occurrence of two distinct patterns.

First, an alteration has been made to the dimensions of the spikes. Increasing the amount of chloroform results in a progressive reduction in the size of the spikes, with a decrease of almost 90% compared to normal levels observed at the highest tested concentrations. This can be considered as an equivalent to going into sleep, as happens with living substrates.

Second, the inter–spike intervals decrease as the dosage of chloroform increases.

These findings indicate that proteinoid systems exhibit an adaptive response to the chloroform exposure rather than undergoing complete degradation. In the field of biology, adaptations indicate an ability to withstand and recover from disruption. The proteinoids exhibit recovery against disruption up to a certain threshold. Their chemical identity remains stable as evidenced by modulation of spike characteristics. This might be seen as analog of the remarkable ability of living cells to rapidly regenerate structures in response to damage.

By understanding the precise mechanisms by which proteinoids sustain electrical functionality despite damage, we can unveil novel approaches to engineer highly resilient computers by drawing inspiration from distinctive biological concepts. More precisely, we may utilise the same mechanisms that enable proteinoids to adjust their signals in response to chloroform in order to construct systems that can endure disturbance. By emulating the mechanisms via which proteinoids regulate signal intensity, synchronise time, and redirect channels as needed, it might be possible to design computers that ensure uninterrupted transmission of information, even under severe attack.

There has been no recorded direct chemical reaction between chloroform (CHCl_3_) and the dipeptide L–Glu:L–Arg. However, in a biological context, chloroform is a well–known membrane disruptive agent, and it is normally not reactive with amino acid side chains under physiological conditions. However, contact can occur indirectly when the solvent disrupts the folding and secondary structure of proteinoids, including L–Glu:L–Arg. The dominant interaction pathway is through non–covalent, solvent–mediated perturbation, rather than direct covalent modification:

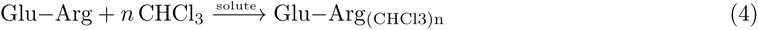

where the binding of *n* chloroform molecules to L–Glu:L–Arg proteinoid aggregates alters the enthalpic state (Δ*H*_solute_) via proteinoid solvation and conformational change.

Rather than reactive chemistry, the impacted secondary structure and folding disrupts control state dynamics:

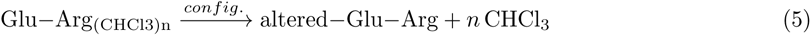

With first order rate dynamics restoring native baseline structures over sub-second timescales after chloroform evacuation.

Figure 17 depicts the influence of chloroform exposure on the electrical characteristics of proteinoid microspheres. The image shows two states: the control proteinoids with prominent spikes (+) before anaesthesia on the left, and the subdued spheres after chloroform treatment on the right. The chloroform molecules, depicted as triangles, are superimposed over the proteinoid population that has been anaesthetised, visually illustrating the inhibitory relationship. The arrows indicate that the electrical muting effect is a result of exposure to chloroform. The text box outlines the significant effects of anaesthesia that have been deduced from experimental findings, including the cessation of neural activity, prevention of external disturbances, restoration of normal function once the anaesthesia wears off, and preservation of the physical structure. This visualisation enhances the stimulus–response data by highlighting the effectiveness of carefully adjusted chloroform in managing previously difficult–to–handle spiking proteinoids for purposes of characterisation and patterning.

The comparative analysis in Tab. 4 shows numerous major contrasts in functional capacities between pure and anaesthetised proteinoids relevant to their use in unconventional computation applications. Pure proteinoids exhibit inherent electrical signalling and associated communication channels, which are disrupted by anaesthesia, leading to a disturbance in both the spontaneous activity and network connectivity. This compromise significantly enhances their ease of manipulation, since anaesthesia allows for direct handling, placement, and integration without being hindered by the need for insulation to protect against signal interference effects. Significantly, the effects of anaesthesia can be reversed, as proteinoids restore their electrical spiking properties once it is removed. The halted spiking behaviours are further supported by the quantitative spike train statistics provided. These recorded variations demonstrate how anaesthesia effectively limits the information capacity of proteinoids, making them more versatile in applications that require flexibility, such as biosensor design and the development of patterned circuits. Additional research can enhance the optimisation of anaesthetic exposure in order to maintain enough processing for certain use scenarios.

**Table 4:** Comparative analysis of computational properties of proteinoids with and without anaesthetic retardation.

| Property | Proteinoids | Anesthetized Proteinoids |
| --- | --- | --- |
| **Electrical Activity** | Spontaneous spikes and oscillations | Suspended spiking, muted signals |
| **Network Connectivity** | Variable inter-proteinoid communication | Disrupted connections |
| **Information Processing** | Complex parallel logic and learning | Restricted processing capability |
| **Handling** | Requires insulation and interference avoidance | Permits direct manipulation and patterning |
| **Reversibility** | Inherent metastable state | Effects reversed upon chloroform removal |

Finally, although anaesthesia allows for the manipulation of proteinoids, the exact mechanism by which it disrupts electrical activity is not well understood. The proposed elements of the chloroform–proteinoid mechanism path are:

- Direct binding of chloroform molecules within hydrophobic pockets on proteinoid surfaces, structurally distorting excitable domains.
- Changes in ion permeability and transport kinetics of membrane altering spike propensity.
- Conformational shifts of intrinsic pore–forming subunits, temporarily blocking conductive states.
- Insertion of chloroform into lipid bilayers modifying membrane dielectric properties and capacitance.
- Combinatorial effects on proteinoid morphology and physiology disrupting synchronized oscillations.

Finally, chloroform, which has been previously used as an anaesthetic, could offer substantial potential in the field of biochemistry and unconventional computing. The agent’s ability to disrupt membranes can modify the folding and secondary structure of proteins, such as proteinoids. This deviation from normal protein behaviour has the potential to be utilised in unconventional computing to influence the processing and storage of information. Proteinoids, which are synthetic proteins that have similar traits to natural proteins, are highly potential contenders for biological computation [68, 69, 70, 4]. Exposure to chloroform causes notable alterations in their electrical characteristics, potentially influencing the development and operation of bio–computational devices. The incorporation of bio–computational modelling of proteinoids interacting with chloroform might be regarded as a distinctive element of biological computing (as seen in Fig. **??**), wherein biological systems regulate or carry out computational processes. Moreover, this specific biochemical interaction, which results in changes in neuromorphic signals and synthesis processes, might potentially be investigated as a subfield within the realm of biological computing, particularly in the context of encoding data.

## 6. Conclusion

By conducting experiments on programmable bio–abiotic substrates, researchers can effectively lead the development of new hardware architectures that have adaptive characteristics. Specifically, by reproducing the observed processes involved in amplitude modulation, time recalibration, and pathway remapping, we can provide a foundation for specialised neuromorphic computers that can maintain stable performance even in the face of external disturbances or attempts to disrupt information. This study uncovers uncharted opportunities within the realm of unconventional computing through an in–depth exploration of the correlation between instability and inventiveness in model biophysical systems. Quantification of spiking potential characteristics shows a progressive decrease in excitation heights and variability in response to sequential chloroform insertion, indicating a dose-dependent modulation of the proteinoid assembly network’s dynamic range. This supports the idea of a balance in these assemblies, as the initial robustness is gradually replaced by deterioration at amplified exposure levels like 25 mg/L, suggesting an optimisation pathway based on a balance between disorder induction and order reconstruction. The considerable reduction in the intervals between spikes suggests increased processing speed, maybe due to chloroform infiltration–induced swelling instabilities. The network’s resilience in retaining coherency despite external perturbations reveals that proteinoid systems may adapt to changes and may promote new organisational patterns. Our findings can help direct engineering pursuits towards the development of hybridized biological–synthetic processors, uniquely designed for customised adaptation processes.

## Data Availability

The datasets generated and analyzed during the current study are available in the zenodo repository, which can be accessed through the following web address: [https://zenodo.org/records/10443058].

## Acknowledgement

The research was supported by EPSRC Grant EP/W010887/1 “Computing with proteinoids”. Authors are grateful to David Paton for helping with SEM imaging and to Neil Phillips for helping with instruments.

## 7. Appendix

The appendix presents quantitative evaluations of the shifts in amplitude and frequency seen in the electric activity of the proteinoid–chloroform system.

**Figure 18:**
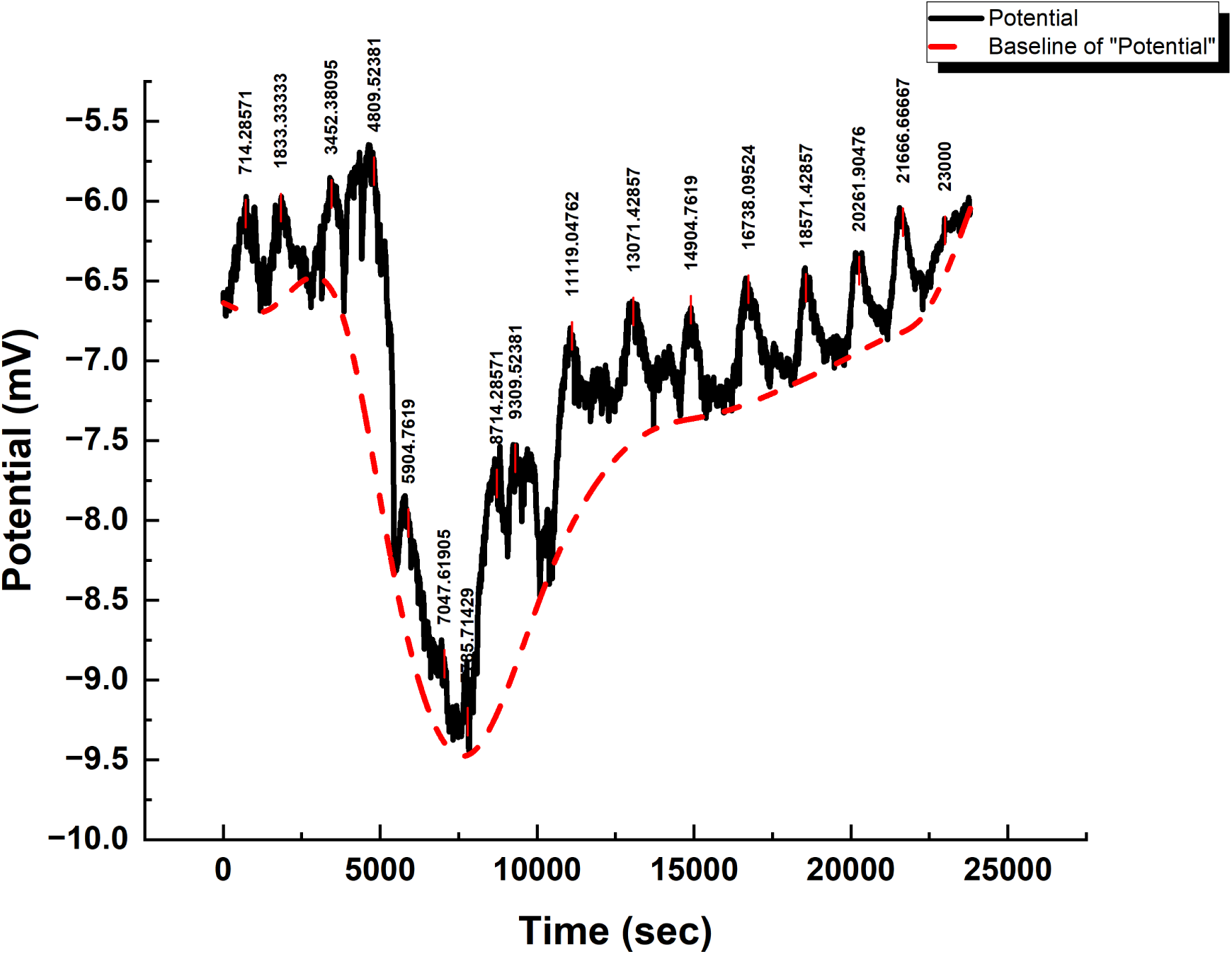
A typical potential signal for a control proteinoid sample (L–Glu:L–Arg). The rhythm of oscillations throughout time is depicted, with a mean periodicity determined from spike analysis of 1392.86 s. The amplitudes range from 0.36 mV to 1.91 mV, with an average of 0.89 mV (mean standard deviation). The positive skewness (1.37), as well as the high kurtosis (3.91), indicate an uneven distribution of spike amplitudes weighted towards more extreme intensity deviations. Even at control settings without external stimulation, the fluctuation in amplitude and timing of oscillations highlights complicated excitation dynamics.

**Figure 19:**
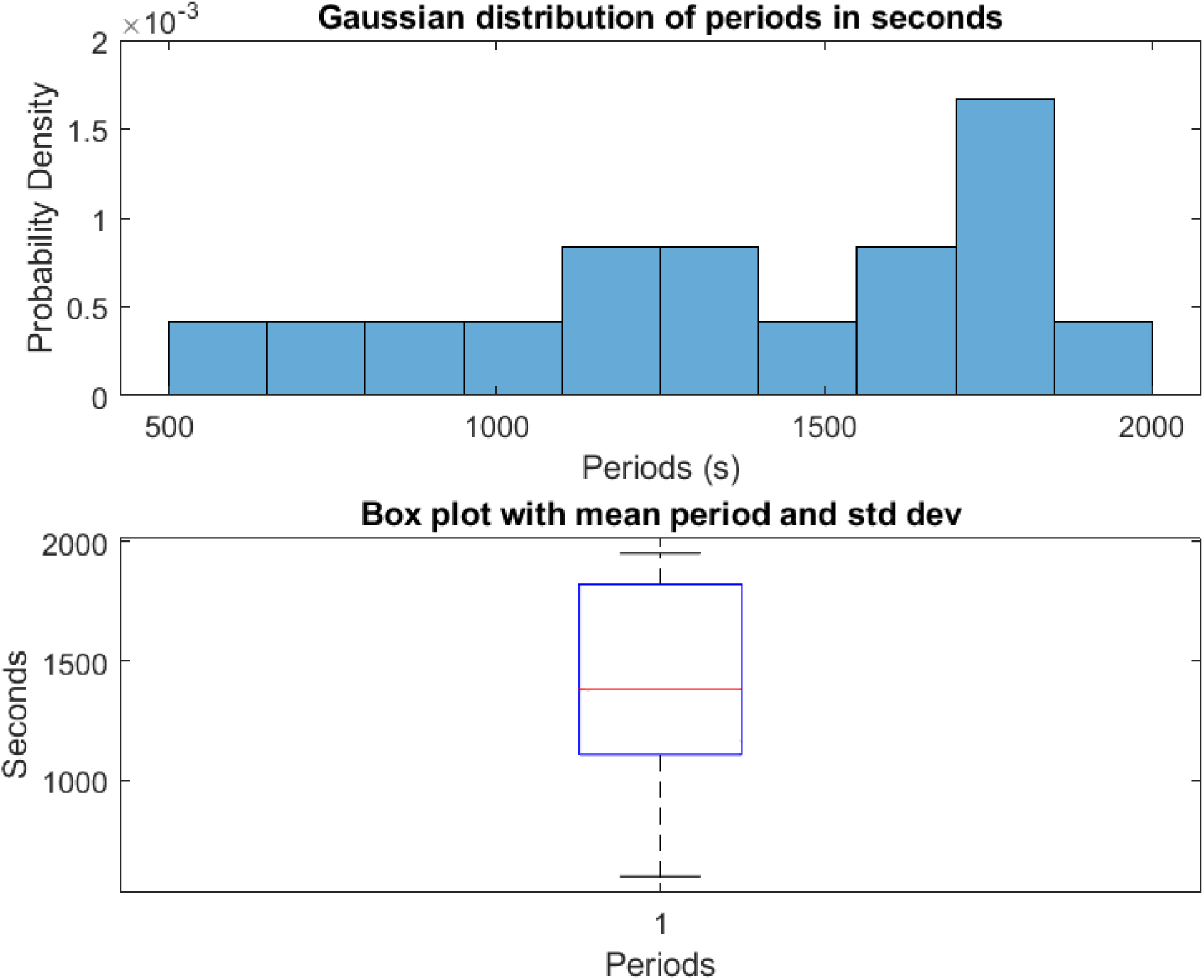
Quantification of spiking period dynamics for L–Glu:L–Arg proteinoid samples at control conditions. (A) Histogram of measured periods fitted to a Gaussian distribution (*µ* =1392.9 s,*σ* = 425.8 s). (B) Boxplot emphasising control period summary data such as mean, standard deviation, minimum and maximum values observed (mean = 1392.86*±*106.46 s; min = 595.24 s; max = 1952.38 s). In the absence of chloroform, the distributions indicate the variability and range of proteinoid interspike duration. Kurtosis of 1.95 and skewness of –0.36 further characterise the shape. Analysing parameters that influence baseline proteinoid excitation tempo may reveal processes that drive emergent oscillatory behaviours. Changes in response to external stimuli may reveal modulatory effects.

**Figure 20:**
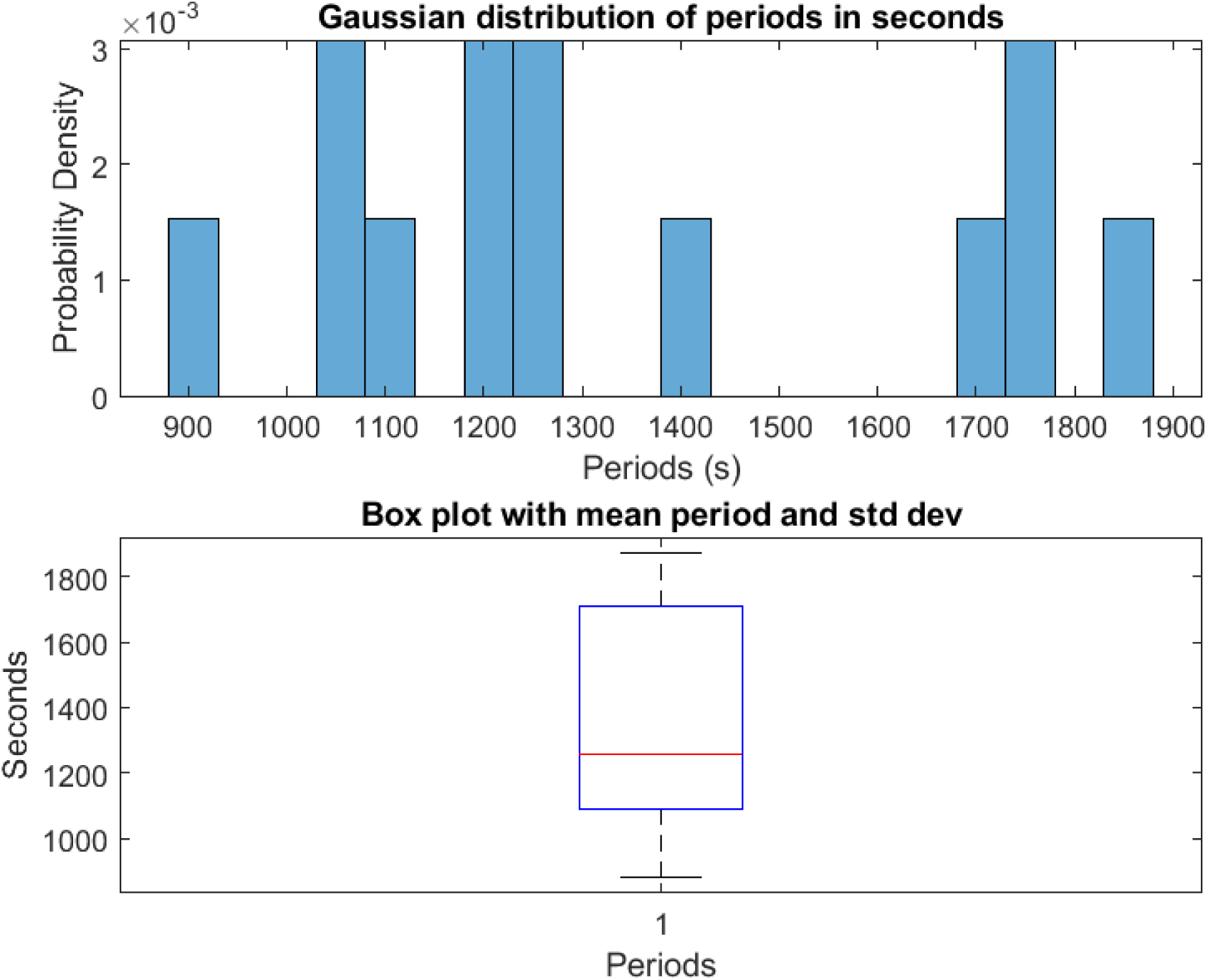
Interspike period dynamics of proteinoid assemblies exposed to low levels of chloroform vapour. The introduction of chloroform vapours from a localised 0.5*×*0.5 cm^2^ filter paper resulted in a skewed distribution of spiking periods relative to the control (mean = 1344.85*±*89.18 s; range 882–1871 s). The periods had a lower standard deviation (321.53 s) compared to the baseline, as well as a lower kurtosis (1.82) and a higher positive skeweness (0.41). Gradual increases in chloroform levels may further influence dynamics. Comparing excitability, conductance, and morphological changes in synthetic protein polymers to external chemical stimuli throughout exposure regimes can help to elucidate adaptive response mechanisms.

**Figure 21:**
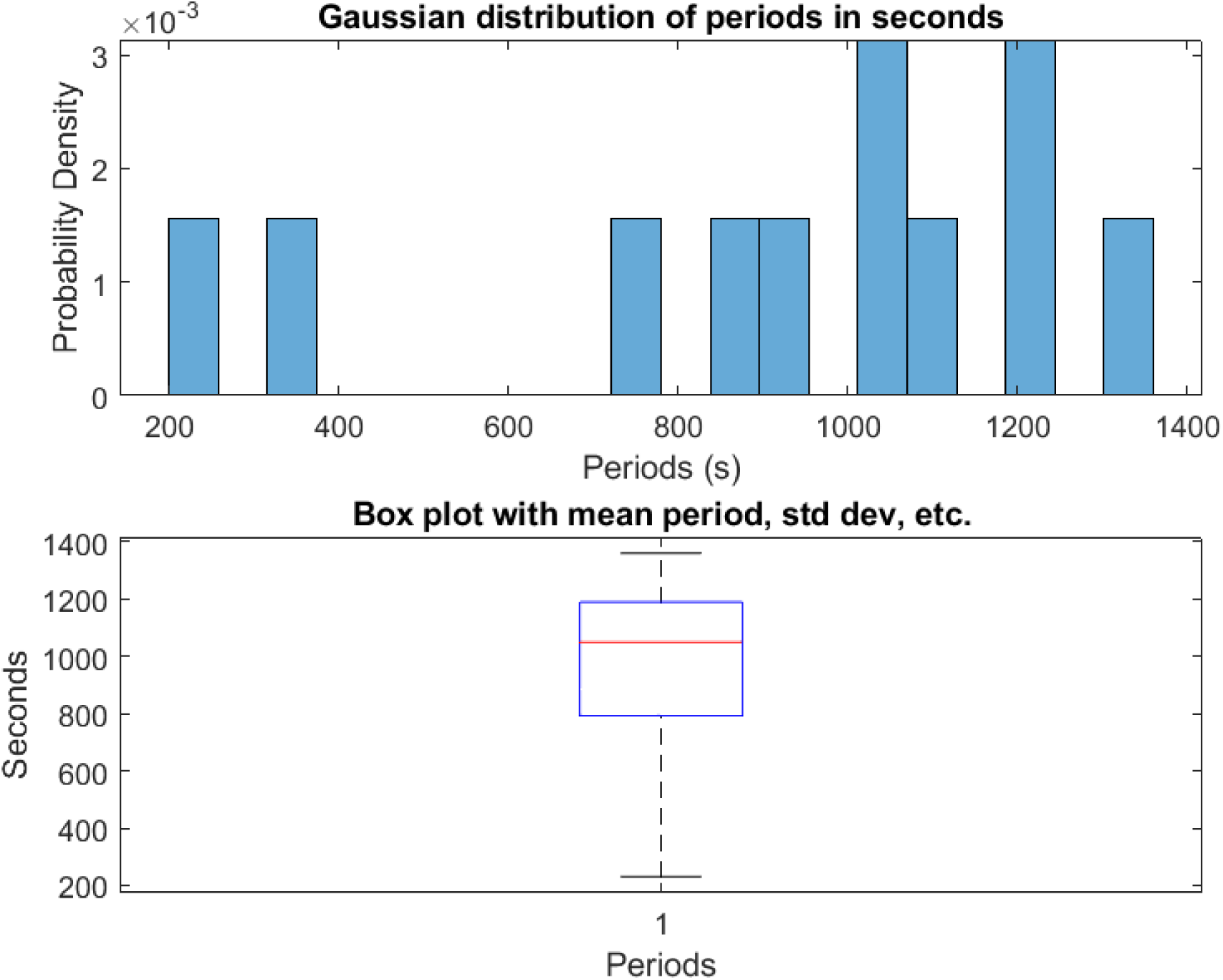
Distribution and statistics of proteinoid inter–spike periods upon sustained chloroform exposure. (A) Periods histogram exhibiting Gaussian traits (mean = 923.00 s; standard deviation = 356.92 s). (B) Boxplot highlighting considerable interval shortening (mean = 923.00*±*107.62 s) compared to higher baseline values without chloroform. The new distribution shows negative skew (skewness = −0.87) with lowered kurtosis (2.67). The considerable leftward shift in periods substantiates substantially greater proteinoid sample excitability.

**Figure 22:**
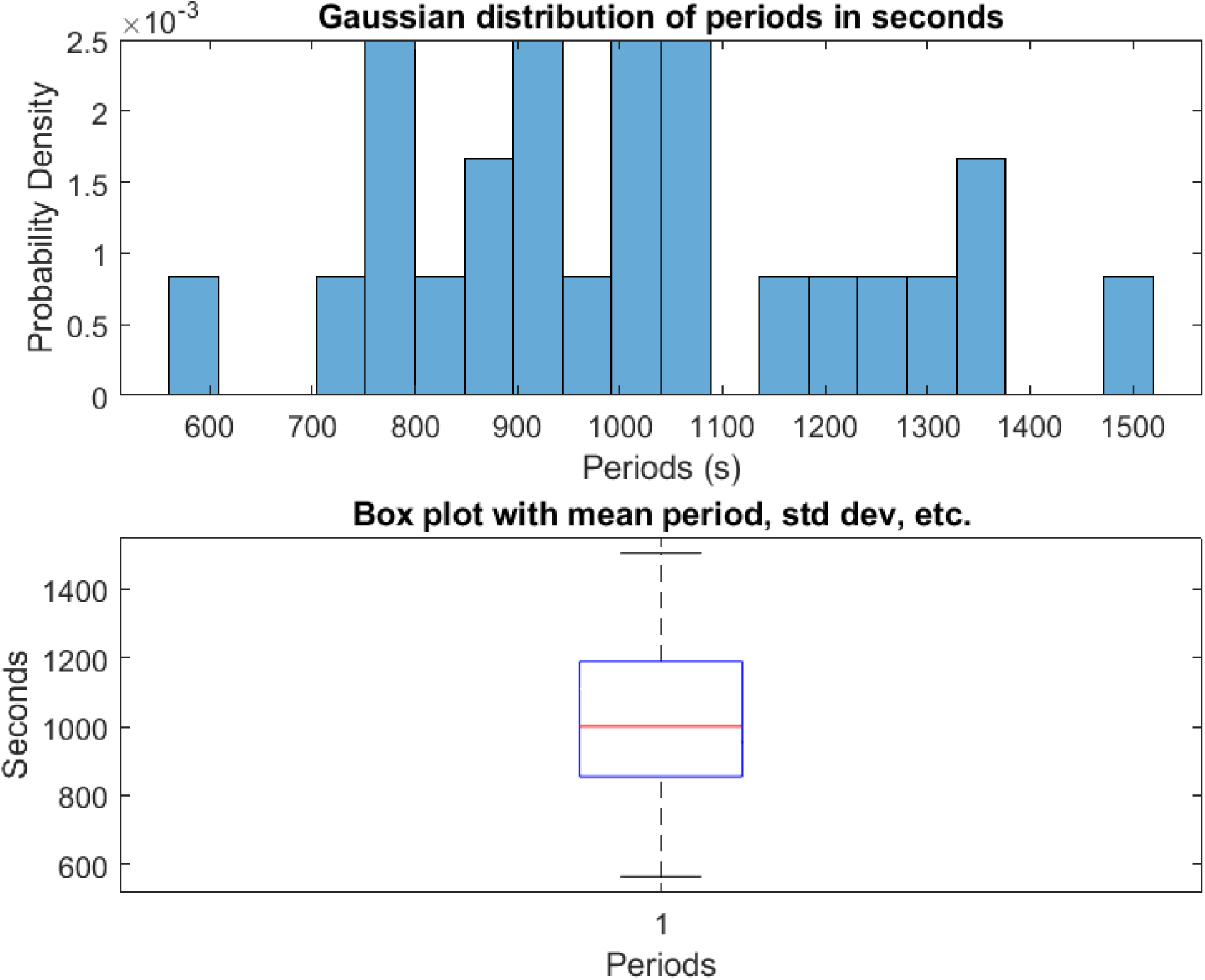
Distribution and statistics of proteinoid inter–spike periods with 3*×*3 cm^2^ chloroform exposure. (a) Histogram of measured periods between spikes fit to a Gaussian distribution (mean = 1011.84*±*45.60 s). (b) Boxplot highlighting the observed range from minimum 563.00 s to maximum 1505.00 s (standard deviation = 228.01 s). The periodicity exhibits a right–skewed distribution (skewness = 0.31) with kurtosis of 2.54. While representing further interval shortening compared to baseline, robust spiking rhythms continue under these exposure levels.

**Figure 23:**
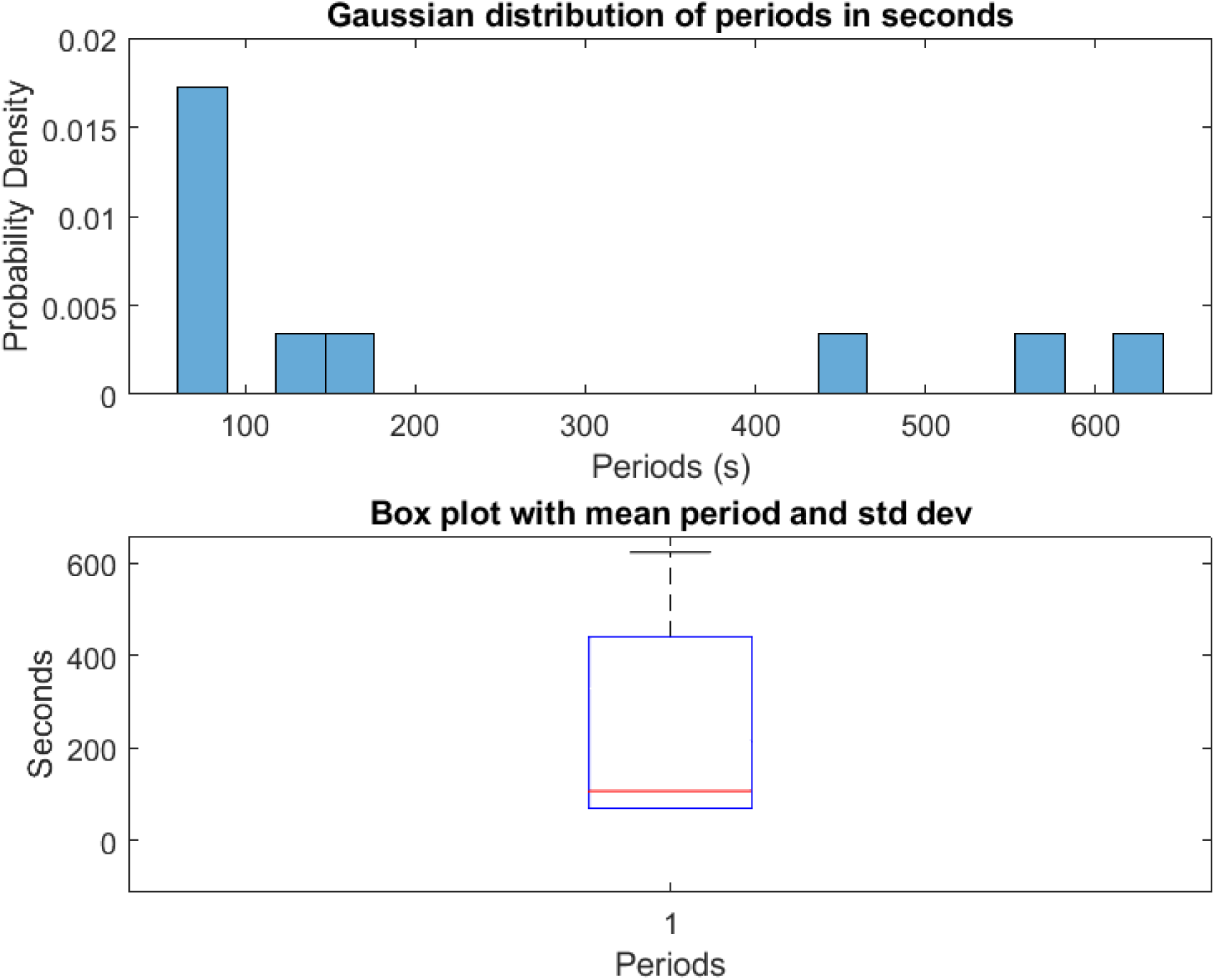
The dynamics of the inter–spike period for proteinoids exposed directly to 25 mg/L chloroform. (A) A periods histogram with reduced intervals that conform to a Gaussian distribution (standard deviation = 223.91 s) (mean = 228.20 s). (B) Boxplot illustrating significant reduction in intervals (maximum period = 623.00 s; minimum period = 69.80 s), exhibiting a positive skew (0.92) and a decreased kurtosis (2.08). Incorporating chloroform directly into proteinoids increases their excitability; rapid, variable spike rates indicate an increase in ion flow. Gaining insight into the mechanisms that govern acute electrochemical sensitivity to chloroform and delineating dose–timed responses will provide clarity regarding the tuning principles that govern solvent–gated bio–electronic materials.

**Figure 24:**
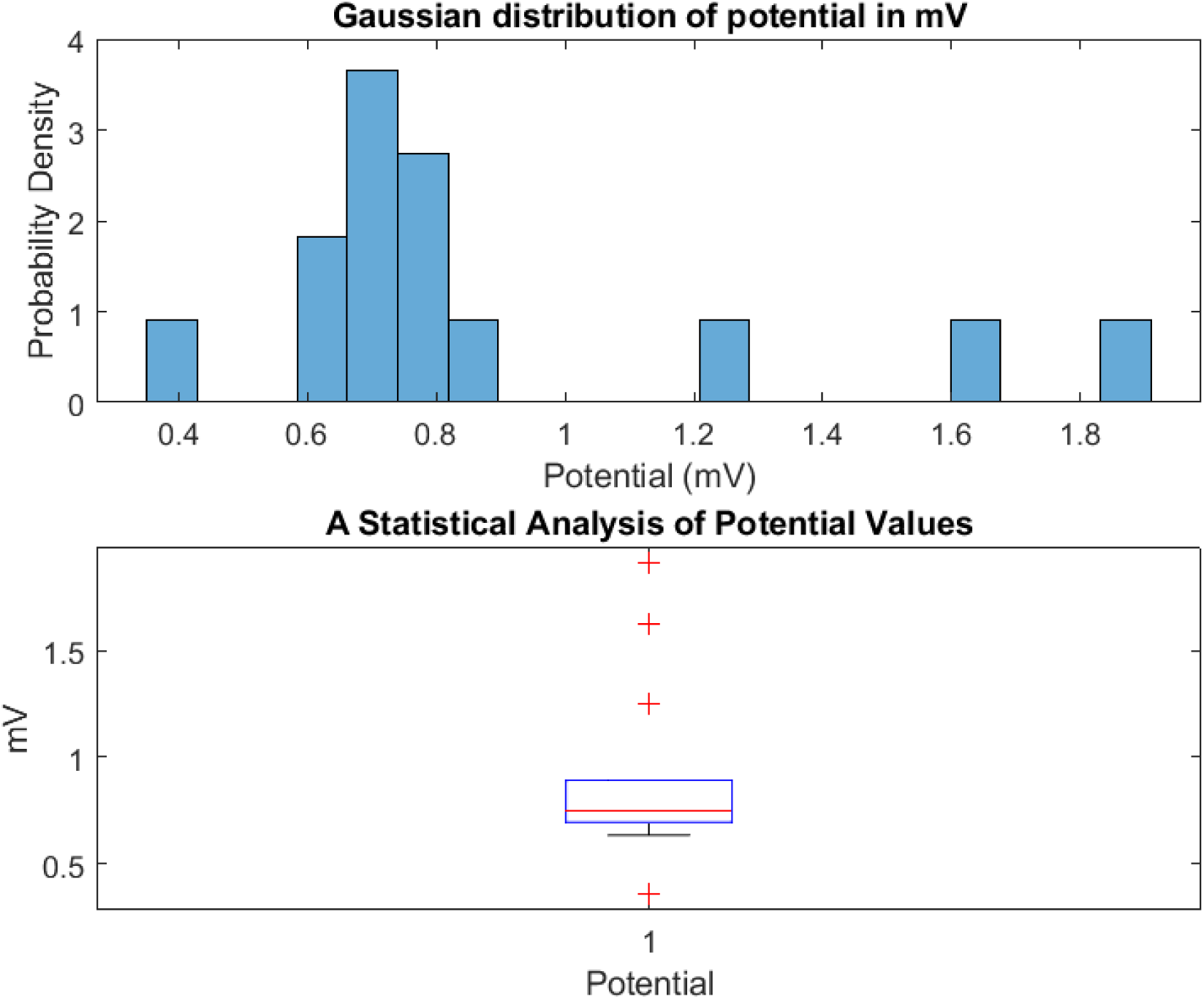
Distribution and statistics of control proteinoid spiking potentials. (A) Spike amplitude histogram exhibiting Gaussian traits (mean = 0.895 mV; standard deviation = 0.419 mV). (B) Boxplot of summary metrics including elevated mean, higher variability, and extended range (mean = 0.895*±*0.112 mV; max = 1.909 mV; min = 0.361 mV). The positively skewed distribution (skewness = 1.37) with high kurtosis (3.91) reflects heterogeneous excitation dynamics in the native proteinoid assemblies.

**Figure 25:**
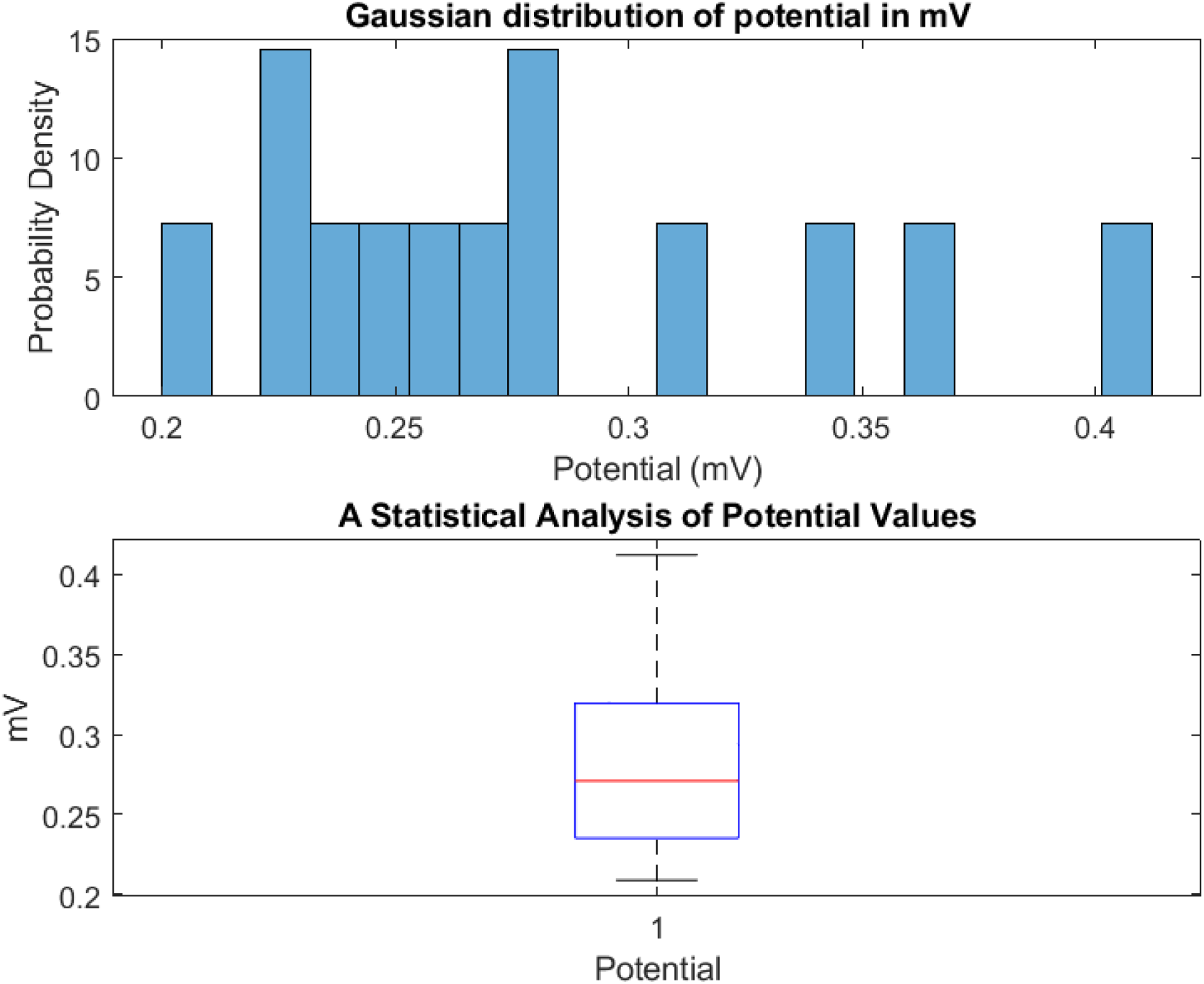
Distribution and statistics of proteinoid spiking potentials upon localized chloroform exposure (0.5*×*0.5 cm^2^. (A) Histogram of measured spike potentials fit to a Gaussian curve (mean = 0.281 mV; standard deviation = 0.059 mV). (B) Boxplot highlighting summary statistics including mean, standard deviation, minimum, and maximum values (mean = 0.281*±*0.016 mV; max spike = 0.412 mV; min spike = 0.209 mV). Introduction of a 0.5*×*0.5 cm^2^ chloroform source induces a considerable downward shift in average proteinoid spike intensity accompanied by distributional changes, yet retains skewed trajectories (skewness = 0.87) and kurtosis = 2.83, indicating preserved bioelectric functionality.

**Figure 26:**
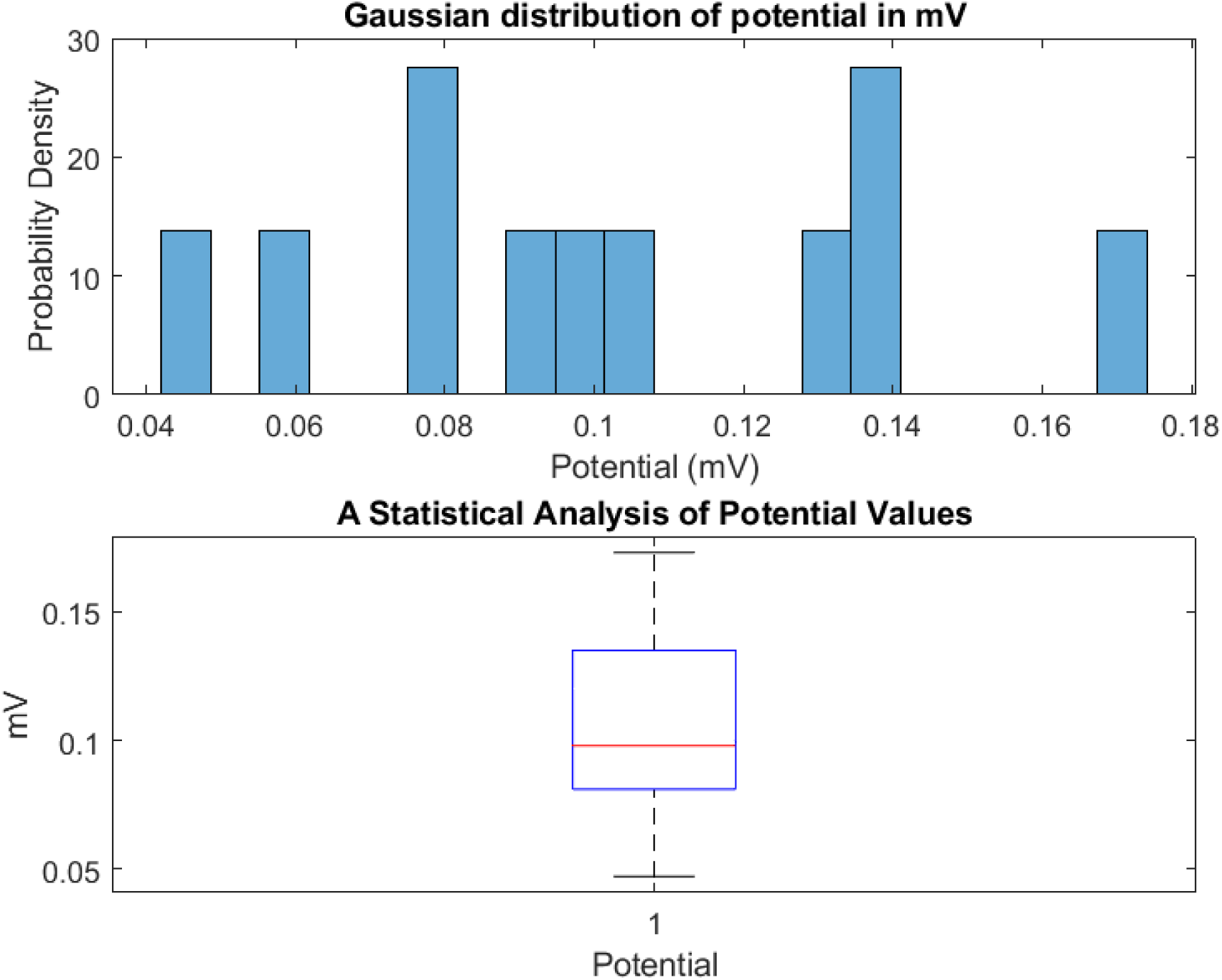
With a 1*×*1 cm^2^ chloroform exposure, the amplitude of proteinoid spiking adjusts. (A) Attenuation of amplitude over time. (B) Box plot of lower amplitudes (mean = 0.104*±*0.011 mV; SD = 0.037 mV) in comparison to higher baseline values in the absence of chloroform. Positive skew is maintained (skewness = 0.31), but kurtosis is reduced (2.26), with a maximum spike of 0.173 mV. The persistence of amplitude modulation and confined distributions suggests increasing dose–dependent response dynamics, whereas the preservation of non–Gaussian bioelectric characteristics indicates persistent excitability.

**Figure 27:**
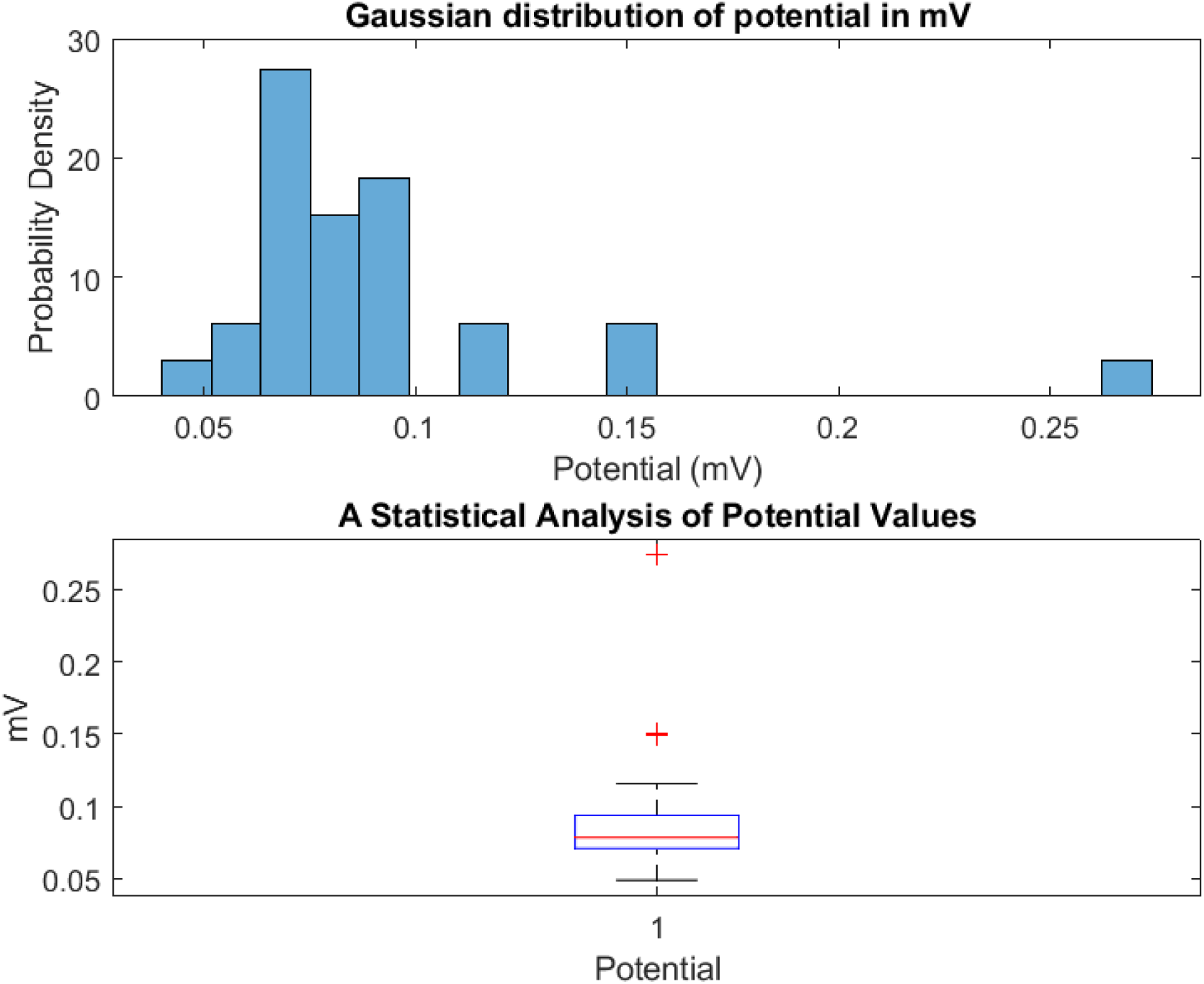
Analysis of the distribution and statistical characteristics of proteinoid spiking potential following exposure to a 3*×*3 cm^2^ area treated with chloroform. The spike amplitude measurements in the histogram show a positive skewness of 2.99, indicating a longer tail on the right side of the distribution. The histogram also exhibits leptokurtic qualities with a kurtosis of 12.93, indicating a higher peak and heavier tails compared to a normal distribution. These traits are induced by transient high intensity outliers. The Gaussian curve fitting analysis reveals that the mean spike potential is 0.091*±*0.008 mV, which is attenuated under the given conditions. (b) The boxplot shows a significant decrease in the range of amplitudes (maximum = 0.274 mV; minimum = 0.049 mV) and a decrease in standard deviation (0.043 mV) compared to higher control values in the absence of chloroform. Examining adaptations across different levels of exposure provides insight into how concentration– dependent response dynamics and processes contribute to non–linear conducting behaviours in disrupted proteinoid systems.

**Figure 28:**
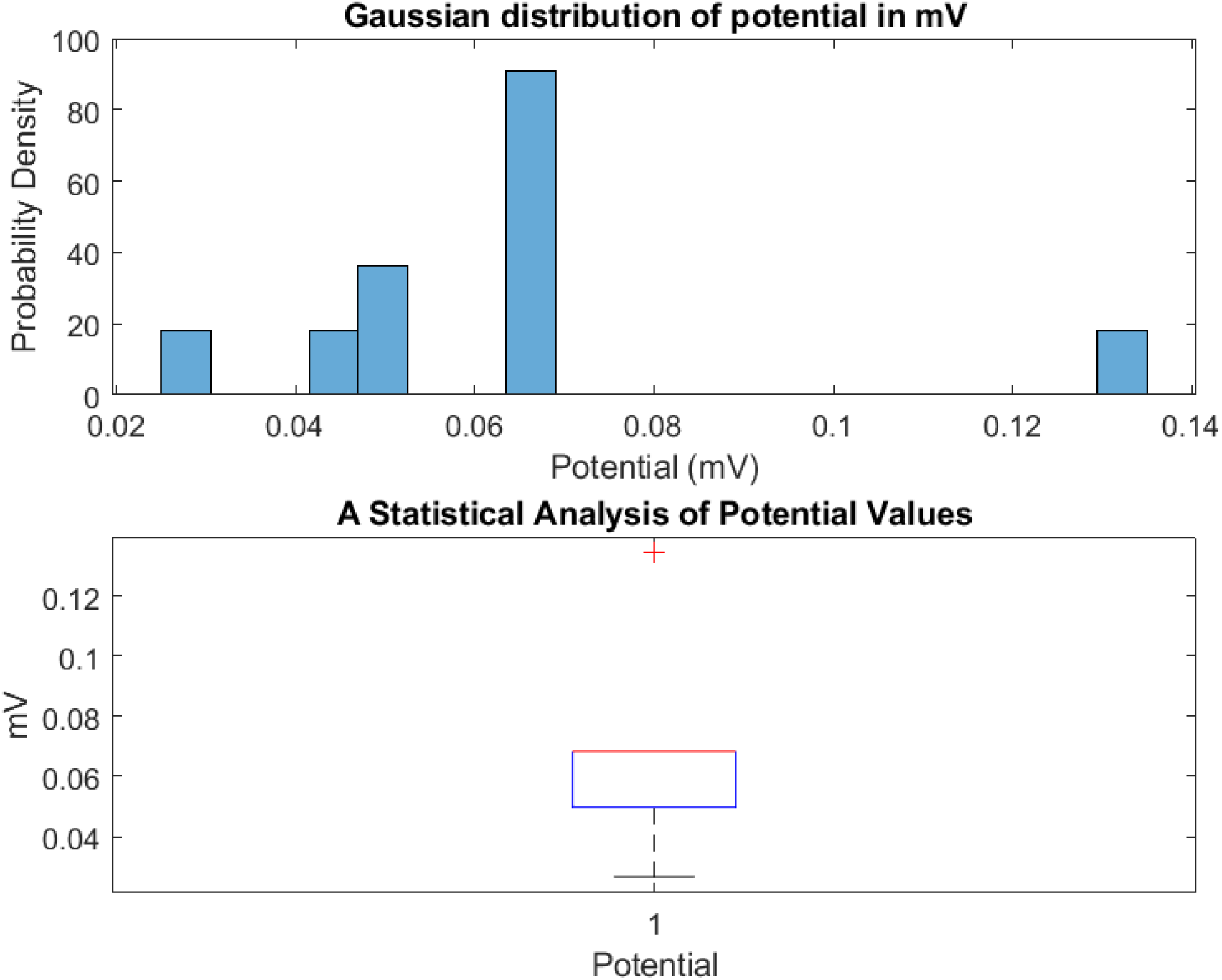
Amplitude and distribution characteristics of proteinoids diluted with 25 mg/L chloroform. (a) The intensity histogram displays a significant mean attenuation of 0.009 mV (0.065*±*1.39 mV), which aligns with a leptokurtic distribution (kurtosis = 5.05) that is positively skewed (skewness = 1.39; standard deviation = 0.028 mV). (b) A boxplot highlighting extreme amplitude range reductions (maximum spike of 0.134 mV; minimum spike of 0.027 mV). Maintaining waveforms and non–Gaussian characteristics indicate persistent self–organized conduction, albeit considerably attenuated, in the presence of a direct solvent conduct.

**Figure 29:**
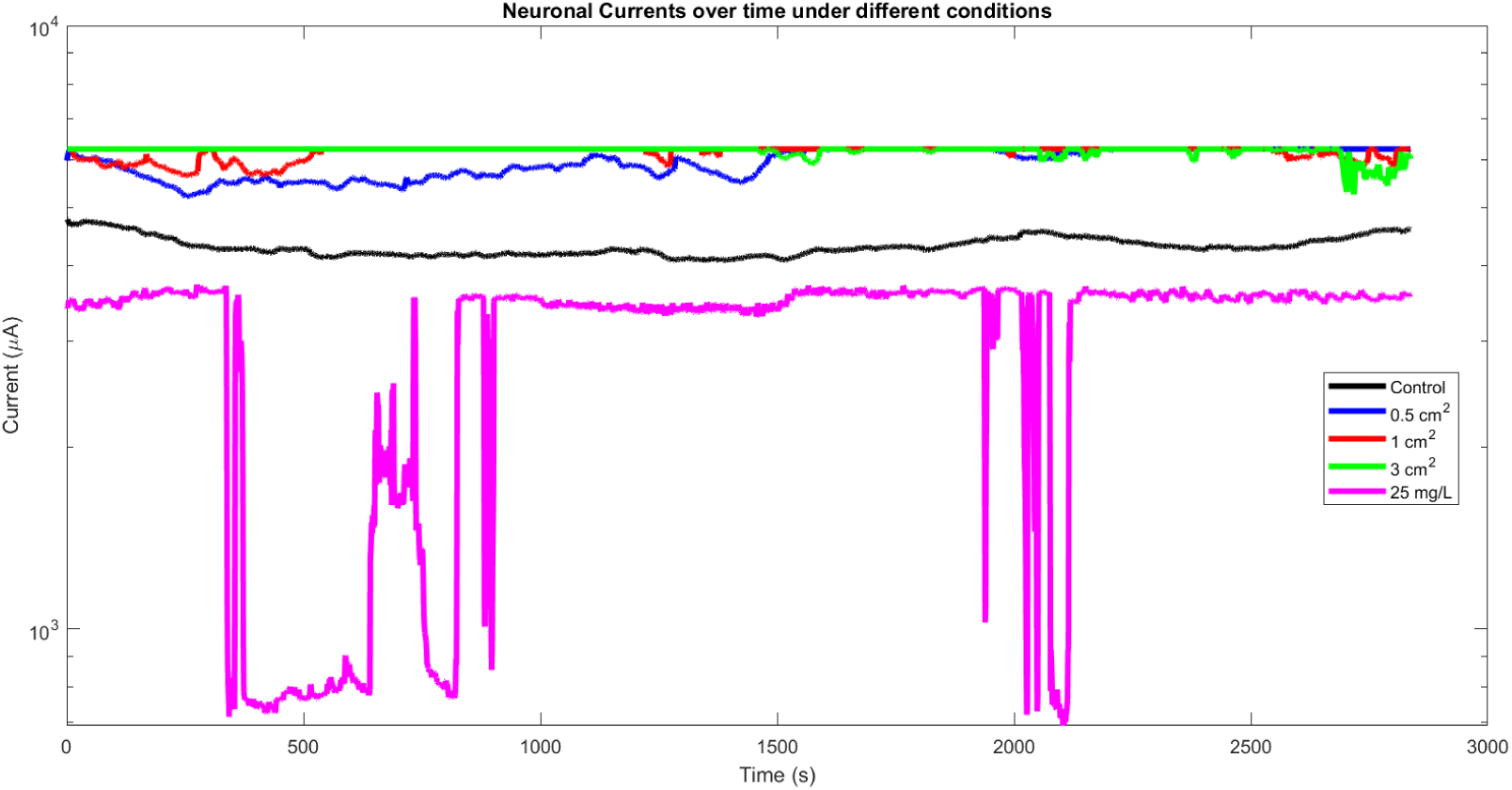
Logarithmic comparison of neuronal currents over time for different proteinoid conditions. The plot exhibits the temporal evolution of neuronal currents in different proteinoids when exposed to chloroform under controlled conditions (“Control”), and in varying surface areas (0.5 cm^2^, 1 cm^2^, 3 cm^2^) and concentrations (25 mg/L). The time is shown in seconds and the currents are presented on a logarithmic scale to better visualise the changes over time. Noticeable differences in the behaviour of the currents highlight the variable effects of Chloroform on the proteinoids.

